# The SiaABC threonine phosphorylation pathway controls biofilm formation in response to carbon availability in *Pseudomonas aeruginosa*

**DOI:** 10.1101/674879

**Authors:** Wee-Han Poh, Jianqing Lin, Brendan Colley, Nicolai Müller, Boon Chong Goh, David Schleheck, Abbas El Sahili, Andreas Marquardt, Yang Liang, Staffan Kjelleberg, Julien Lescar, Scott A. Rice, Janosch Klebensberger

## Abstract

The critical role of bacterial biofilms in chronic human infections calls for novel anti-biofilm strategies targeting the regulation of biofilm development. However, the regulation of biofilm development is very complex and can include multiple, highly interconnected signal transduction/response pathways, which are incompletely understood. We demonstrated previously that in the opportunistic, human pathogen *P. aeruginosa*, the PP2C-like protein phosphatase SiaA and the di-guanylate cyclase SiaD control the formation of macroscopic cellular aggregates, a type of suspended biofilms, in response to surfactant stress. In this study, we demonstrate that the SiaABC proteins represent a signal response pathway that functions through a partner switch mechanism to control biofilm formation. We also demonstrate that SiaABCD functionality is dependent on carbon substrate availability for a variety of substrates, and that upon carbon starvation, SiaB mutants show impaired dispersal, in particular with the primary fermentation product ethanol. This suggests that carbon availability is at least one of the key environmental cues integrated by the SiaABCD system. Further, our biochemical, physiological and crystallographic data reveals that the phosphatase SiaA and its kinase counterpart SiaB balance the phosphorylation status of their target protein SiaC at threonine 68 (T68). Crystallographic analysis of the SiaA-PP2C domain shows that SiaA is present as a dimer. Dynamic modelling of SiaA with SiaC suggested that SiaA interacts strongly with phosphorylated SiaC and dissociates rapidly upon dephosphorylation of SiaC. Further, we show that the known phosphatase inhibitor fumonisin inhibits SiaA mediated phosphatase activity *in vitro*. In conclusion, the present work improves our understanding of how *P. aeuruginosa* integrates specific environmental conditions, such as carbon availability and surfactant stress, to regulate cellular aggregation and biofilm formation. With the biochemical and structural characterization of SiaA, initial data on the catalytic inhibition of SiaA, and the interaction between SiaA and SiaC, our study identifies promising targets for the development of biofilm-interference drugs to combat infections of this aggressive opportunistic pathogen.

**Author Summary:** *Pseudomonas aeruginosa* is a Gram-negative bacterium that is feared within clinical environments due to its potential to cause life-threatening acute and chronic infections. One cornerstone of its success is the ability to form and disperse from biofilms, which are self-made, multicellular structures that protect the individual cell from the human immune system and antibiotic treatment. As such, therapies that combine a biofilm-interference strategy and the use of antimicrobial drugs represent one of the promising strategies to tackle infections of this organism. With the current study, we gain a deeper understanding of the SiaABCD mediated biofilm formation in response to clinically relevant environmental conditions. Further, our structural and biochemical characterization of the PP2C-type protein-phosphatase SiaA and the partner switch protein SiaC suggest that both represent promising novel targets for the development of future anti-biofilms drugs based on a signal interference strategy.

## Introduction

Biofilms are multicellular structures embedded in a self-made matrix, which can occur attached at a solid surface, at the gas-liquid interface as a pellicle, or freely floating as suspended aggregates (flocs) in the liquid phase [1–3]. The ability to form and disperse from biofilms is a ubiquitous feature of microorganisms, which includes the genetically controlled production of extracellular polymeric substances (EPS) such as polysaccharides, eDNA and proteinaceous surface adhesins [4–6]. Biofilm formation can be triggered in response to oxidative and nitrosative stress [7,8] and toxic compounds such as antibiotics [9,10], surfactants [11,12] or primary fermentation products [13,14]. Cells embedded in biofilms show increased resistance towards the presence of such stressors compared to their single-cell, planktonic counterparts, which explains the evolutionary success of the biofilm lifestyle [15–19]. As a consequence, biofilms are problematic in health-care settings despite stringent hygiene regimes [20–22]. Biofilms are also associated with the establishment of chronic infections that are recalcitrant to the effects of the host immune system and therapeutic interventions [23–27] and hence are almost impossible to eradicate with conventional therapies [28–31].

The regulation of biofilm development and virulence traits is complex and can include multiple, highly interconnected signal transduction pathways [6,32–36]. In addition to quorum sensing (QS) [37,38], nucleotide-based second messengers and changes in protein phosphorylation represent key mechanisms for the regulation of these physiological changes [39–42]. Protein phosphorylation is most often mediated by two-component systems (TCSs) or chemosensory signalling pathways [43–45]. TCSs are typically composed of a sensor kinase and a cognate response regulator that elicits a specific response upon its phosphorylation. Most sensor kinases in bacteria belong to the family of histidine kinases, transferring a phosphoryl group from a conserved histidine of its transmitter domain to a specific aspartate residue in the receiver domain of the corresponding response regulator. This aspartate phosphorylation is labile and, thus, specific phosphatases are usually not required to inactivate the response regulator over time. In contrast, the phosphorylation of a serine or threonine residue by serine/threonine kinases (STK), are more stable and thus require the additional presence of a protein phosphatase, e.g. protein-phosphatase family-2C (PP2C)-like phosphatases, to facilitate a reversible regulation [46,47]. *Pseudomonas aeruginosa*, an opportunistic human pathogen of critical concern, is an ubiquitous organism that can thrive in multiple environmental niches and genetic evidence indicates that infections usually arise from environmental sources [48–51]. Various regulatory pathways affect its biofilm formation and virulence traits, including, but not limited to, the GacS/GacA system, the threonine phosphorylation pathway (TPP) and the Wsp, Yfie or HptB pathways [24,52–57]. While *P. aeruginosa* does not generally infect healthy humans, it is a serious threat in hospital environments, particularly for individuals with burn wounds or chronic diseases such as obstructive pulmonary disease (COPD) and the genetic disorder cystic fibrosis (CF)[28,58–62]. In addition to surface attached biofilms, *P. aeruginosa* also forms suspended biofilms (i.e. cellular aggregates)[63,64], which are medically relevant as they are regularly observed in chronic infections [26,27,65–67]. Moreover, suspended biofilms have been shown to greatly influence the development, structure and function of their surface-attached counterparts and have thus been suggested to play an important role in niche colonisation and chronic infections [68,69].

We previously demonstrated that PP2C-like protein phosphatase SiaA and the di-guanylate cyclase (DGC) SiaD regulate the formation of large, macroscopic, DNA-containing suspended biofilms in response to the surfactant sodium dodecyl sulfate (SDS) in a c-di-GMP-dependent manner [11,70–72]. We now report that all genes encoded in the *siaABCD* operon are involved in regulation of the cellular aggregation in response to carbon availability with a variety of growth substrates. Our study demonstrates that the phosphorylation status of the SiaC protein at threonine 68 (T68) is crucial for this regulation, and that phosphorylated SiaC represents a substrate for the phosphatase SiaA and its kinase counterpart SiaB, which is in line with results of a recently reported study [73]. In addition, the crystal structures of the PP2C-domain of SiaA and of the SiaC protein provided insights into their catalytic and regulatory mechanisms.

## Results

### SiaABCD is involved in biofilm formation during growth on SDS and glucose

Cells of *P. aeruginosa* preferentially grow as macroscopic suspended biofilms (cellular aggregates in the mm range) in minimal medium with SDS [11]. To investigate whether all genes of the *siaABCD* operon [70] are involved in biofilm formation under these conditions, we tested individual mutant strains in microtiter plates for macroscopic aggregation and compared it to cultures growing on glucose (**Fig 1**).

**Fig 1:**
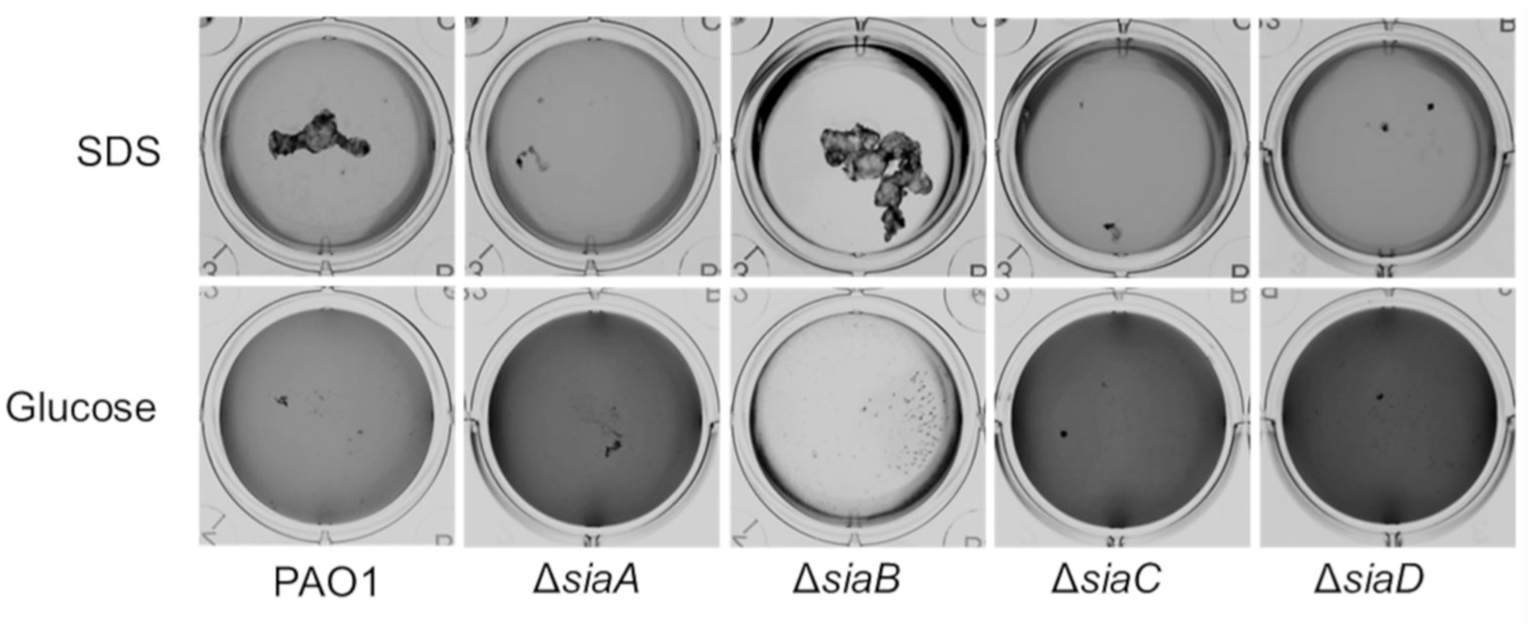
Growth of the *P. aeruginosa* strains on 3.5 mM SDS or 22 mM glucose in 12 well microtiter plates after 18 h, at 30°C and 200 rpm. Images were acquired using a flat-bed scanner (Umax Powerlook), normalized by the “match colour” function of Photoshop (Adobe Photoshop CS5), and converted to greyscale.

As shown previously [70], growth on SDS led to the formation of macroscopic aggregates in the parental strain but not in the Δ*siaA* and Δ*siaD* mutants. Likewise, no macroscopic aggregates were observed in cultures of Δ*siaC*, whereas Δ*siaB* showed increased aggregation. The aggregative behaviour of cells was inversely correlated with the optical density (OD_600_) of the surrounding medium with higher OD_600_ values for strains Δ*siaA*, Δ*siaC* and Δ*siaD* (darker colour in normalized pictures) compared the parental strain or the Δ*siaB* mutant. In contrast, no macroscopic aggregates were found during growth with glucose for all strains. However, the OD_600_ of the culture medium of glucose-grown cultures followed a similar pattern as those grown on SDS. Growth on glucose has previously been found to occur mainly as attached or suspended biofilms (aggregates of up to 400 nm in size), which quickly disperse upon carbon starvation in a c-di-GMP-dependent manner [63]. To study, whether differences in the ability to form suspended biofilms could explain the observed OD_600_ pattern of the *sia*-mutants, we analysed glucose-growing cultures with scanning electron microscopy (SEM) and laser diffraction analysis (LDA).

### SiaABCD regulates the ratio of freely suspended and aggregated cells

SEM imaging (**Fig 2**, 10,000×; left panels) showed the presence of three-dimensional structures composed of densely packed microbial cells that projected above the substratum. The multicellular aggregates appeared to be less structured in the Δ*siaA*, Δ*siaC and* Δ*siaD* mutants compared to the parental strain. In contrast, the Δ*siaB* mutant formed much larger and more densely packed aggregates than the parental strain. Similar differences in aggregate sizes were also noted at lower magnification (1,300×), with the additional observation that aggregates were generally less abundant for the Δ*siaA*, Δ*siaC and* Δ*siaD* mutants.

**Fig 2:**
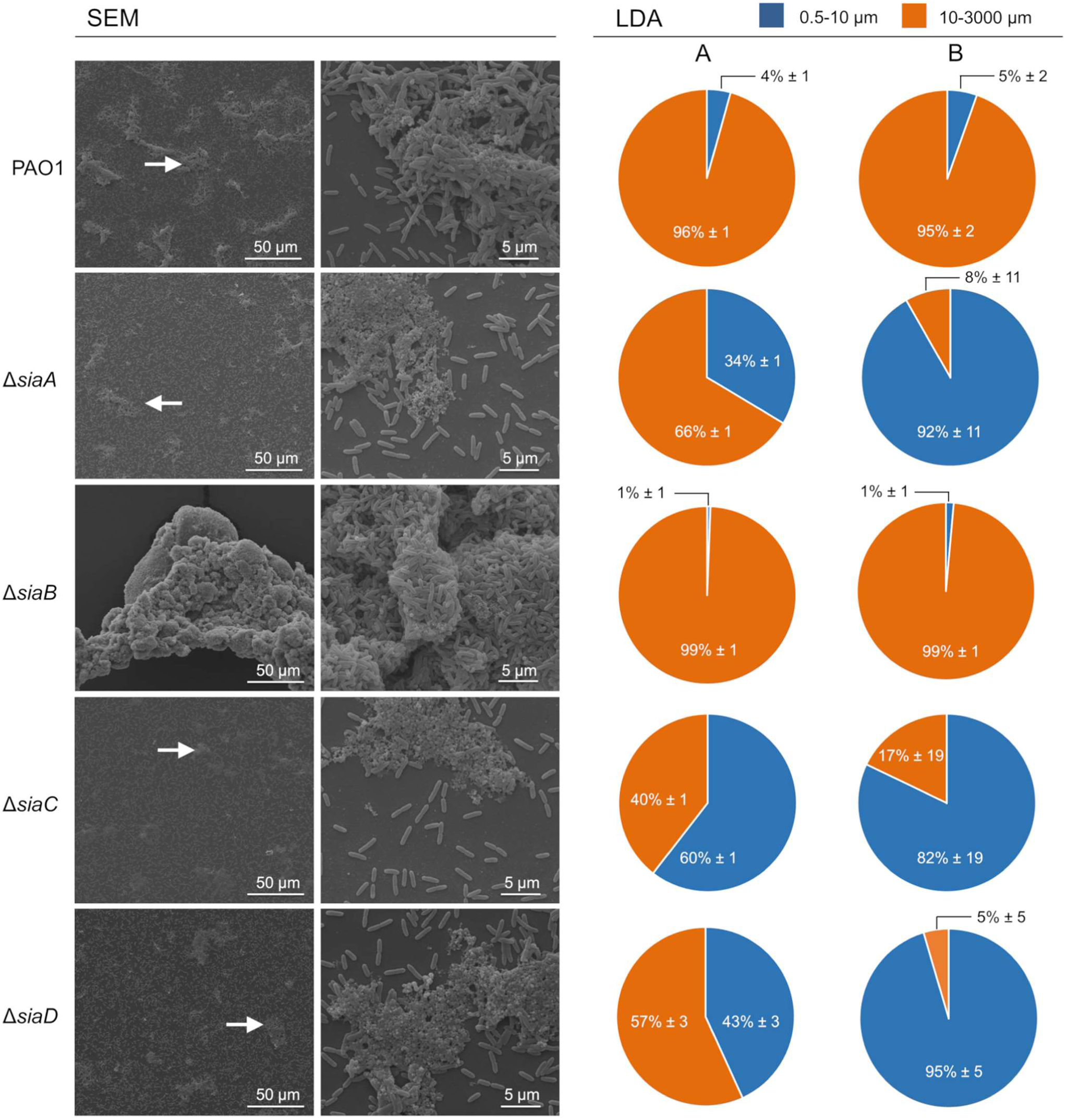
Scanning electron microscopy (SEM) and laser diffraction analysis (LDA) demonstrated the different aggregation phenotypes of *P. aeruginosa* wildtype and mutant strains during growth with glucose. Triplicate cultures of strains were grown with 22 mM glucose in 50 mL cultures shaking at 200 rpm at 30°C. After 5.5 h incubation, the replicate samples were pooled and used for SEM. The white arrows (left panel) highlight representative regions showing aggregated cells, which were then examined at higher magnification (right panel). Similar samples were also analyzed using a particle size analyzer (SALD 3101, Shimadzu; see Materials and Methods for details) in the range 0.5-3000 µm diameter. The data obtained from two independent biological replicates (A and B; n = 2) are presented as the percentages of the total bio-volume of particles distributed over two different size ranges (0.5-10 µm and 10-3000 µm) relative to the total bio-volume of all particles detected. The data represent the mean value of triplicate measurement of the same sample (n = 3) with the corresponding technical error.

Light diffraction analysis (LDA) can be used to determine the size distribution of PAO1 aggregates in liquid cultures^64^. LDA analysis in the current study confirmed these results for cultures of the wildtype strain, which were dominated by particles > 10 µm (≥ 95% ± 2% of the total bio-volume of all particles). Particles < 10 µm, which include the single cells, represented only a minor fraction (≤ 5% ± 2%). In line with SEM results, the distribution of particles in cultures of the Δ*siaA*, Δ*siaC* and Δ*siaD* mutants were strongly shifted towards smaller sizes. A substantial increase in the total bio-volume of all particles < 10 μm for Δ*siaA* (≥ 6.6 fold), Δ*siaC* (≥ 11.8 fold) and Δ*siaD* (≥ 8 fold) was consistently observed. In contrast, for cultures of Δ*siaB* the total bio-volume of all particles > 10 µm (≥ 99% ± 1%) was considerably higher than for the parental strain. Consequently, almost no particles < 10 µm were detected.

### SiaABCD regulates biofilm formation as a response to carbon availability

Biofilm formation is a dynamic process that varies according to the time point used for quantification. The temporal nature of biofilm formation and dispersal can be monitored with a microtiter plate-based biofilm assay [63]. When the impact of the SiaABCD proteins on the biofilm life cycle during growth on glucose was studied using this assay, we found the following temporal pattern of biofilm formation for the wildtype strain: *i)* a steady increase in biofilm formation while glucose was present in excess; *ii)* a decrease in biofilm formation upon exhaustion of glucose; and *iii)* an increase in optical density in the culture supernatant that lagged behind biofilm formation, but continued to increase even after glucose was completely consumed (**Fig 3**). The latter observation is consistent with the dispersal of cells from the attached and/or suspended biofilm into the liquid phase in response to carbon starvation [63]. The Δ*siaA*, Δ*siaC* and Δ*siaD* mutants exhibited a similar growth pattern on glucose as described for the parental strain (**Fig 3** and **Fig S1**). However, all strains showed a decrease in overall biofilm formation and as a result, the increase in OD_600_ upon carbon starvation was also lower. In contrast, the Δ*siaB* mutant predominantly grew as a biofilm with much lower planktonic growth and with a reduced dispersal response upon carbon starvation (percentage in biofilm reduction during consecutive sampling points upon carbon limitation) compared to all other strains.

**Fig 3:**
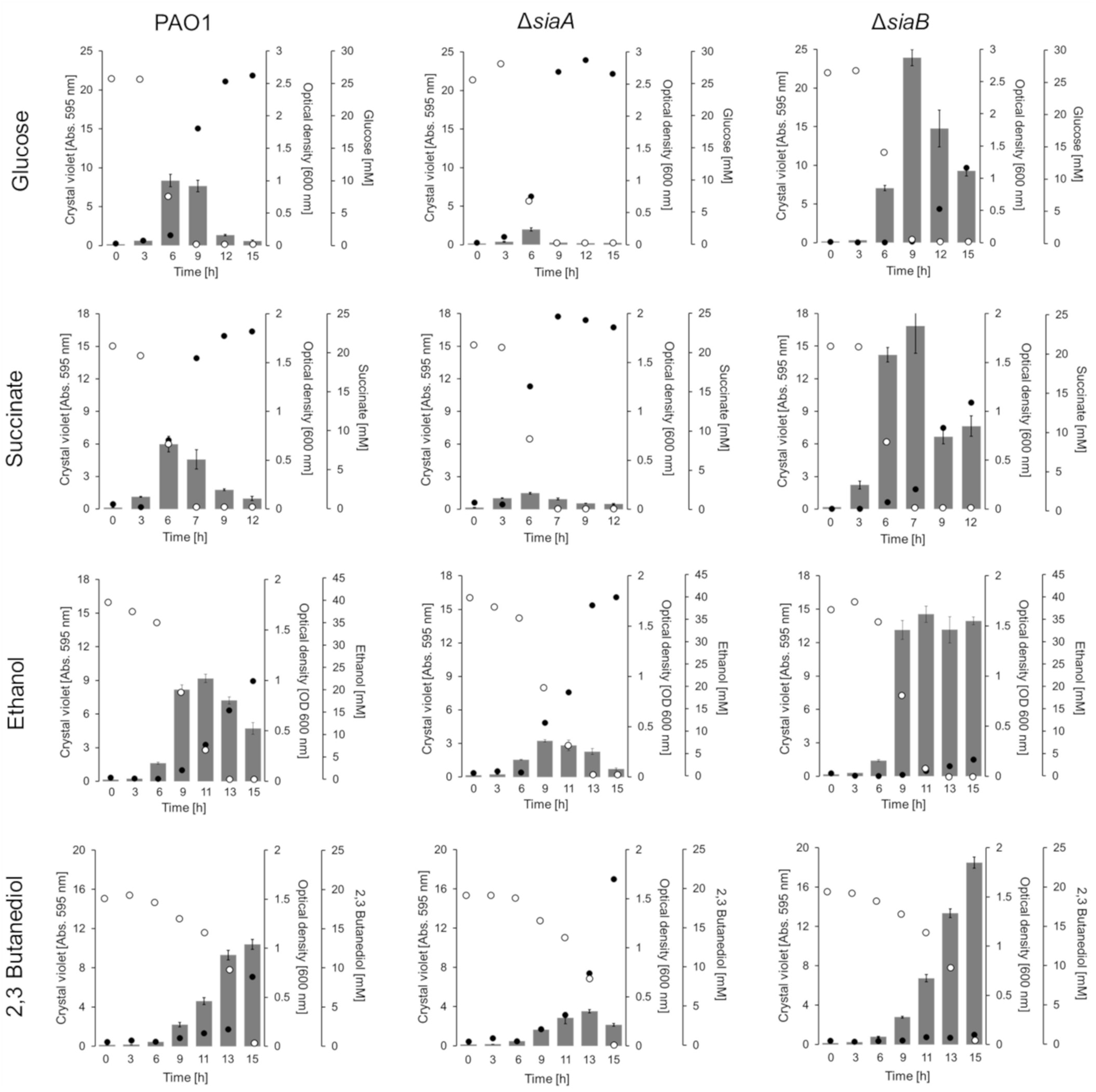
Quantification of biofilm formation and growth in suspension of strains PAO1, Δ*siaA* and ΔsiaB with 22 mM glucose, 20 mM succinate, 40 mM ethanol or 20 mM 2,3-butanediol in 24 well microtiter plates (growth of the Δ*siaC* and Δ*siaD* mutants are shown in **Fig S1**). Individual 24 well microtiter plates were used to quantify attached biofilms by crystal violet (CV) staining (*grey bars*) as well as the growth in the supernatant as OD_600_ (*black dots*); substrate concentrations are shown (*open circles*). For crystal violet (CV) staining, the data are shown as the mean value of four biological replicates (n = 4). To reduce complexity, the standard deviation of replicates is shown as error bars. OD_600_ and substrate concentrations represent the mean value from the same quadruplicates but quantified from pooled (1:1:1:1 [v/v/v/v]) samples using a photometer, the GO assay kit and specific HPLC methods, respectively (see Materials and Methods for details). All cultures were incubated at 30°C and 200 rpm shaking.

To study whether the regulatory impact of the SiaABCD proteins on biofilm formation can be expanded to other carbon sources, we additionally tested succinate, ethanol, 2,3-butanediol (**Fig 3** and **Fig S1)**. For all carbon sources tested, a similar pattern as described for growth on glucose was observed, with the following two exceptions: *i)* no dispersal of the Δ*siaB* biofilm was observed when grown on ethanol upon carbon depletion; and *ii)* no dispersal of the Δ*siaB* and wildtype biofilms was observed during growth on 2,3-butanediol. However, it is important to note that for the later condition, 2,3-butanediol was not completely depleted (≤ 0.6 mM) in the cultures and hence the cells would not have experienced carbon starvation.

### SiaA and SiaB represent a phosphatase/kinase couple that balance the phosphorylation status of SiaC

The SiaA protein is predicted to be a PP2C-like protein phosphatase (PPM-type) (http://www.pseudomonas.com/feature/show/?id=103077&view=functions). A distinctive feature of these phosphatases is their dependency on Mn^2+^ or Mg^2+^ ions as cofactors and serine and/or threonine residues in their target protein (http://www.ebi.ac.uk/interpro/entry/IPR001932). Structure-function predictions of SiaB using I-TASSER suggested that SiaB has protein kinase activity (**Fig 4A**). Notably, SiaC contains a putative phosphorylation site at threonine residue 68 (T68)[74] and as such could represent the target of the predicted protein phosphatase/kinase activities of SiaA and SiaB. To test this hypothesis, we purified the C-terminal part of SiaA (amino acids 386-663; SiaA-PP2C; Genbank: NP_248862) including the putative PP2C-like phosphatase domain (amino acids 453-662**; Fig 4B**) and SiaB protein from *E. coli* by metal-affinity chromatography (**Fig S2**). Similarly, the SiaC protein was purified from *E. coli* or Δ*siaA* lysates and was subsequently analyzed by shotgun peptide-mass spectrometry (PMS; **Table S1**). For the SiaC variant purified from *E. coli*, the T68-containing peptide (peptide [LLYLN**T**SSIK]) was identified predominantly by fragment-masses derived of precursor ions of the parental, non-phosphorylated peptide. In addition, the non-phosphorylated peptide was quantified at > 1000 fold lower intensity (1.4 × 10^11^ vs. 4.0 × 10^14^ peak area) by precursor ions of its phosphorylated analogue; hence, these preparations contained almost exclusively non-phosphorylated protein. In contrast, the SiaC variant purified from Δ*siaA* cell lysates (SiaC^P^), the corresponding phosphorylated peptide was found at a higher abundance than the non-phosphorylated peptide (7.8 x × 10^6^ vs. 3.2 × 10^6^ peak area). Hence, these preparations contained relevant amounts of phosphorylated SiaC^P^ at position T68 (71.27% of total peak area). Incubation of SiaB (0.5 µM) alone or with SiaC^P^ (5 µM) did not consume ATP, whereas incubation of SiaB with SiaC (5 µM) consumed 4.45 µM ATP, indicating kinase activity of the SiaB protein in the presence of SiaC (**Fig 4C**).

**Fig 4:**
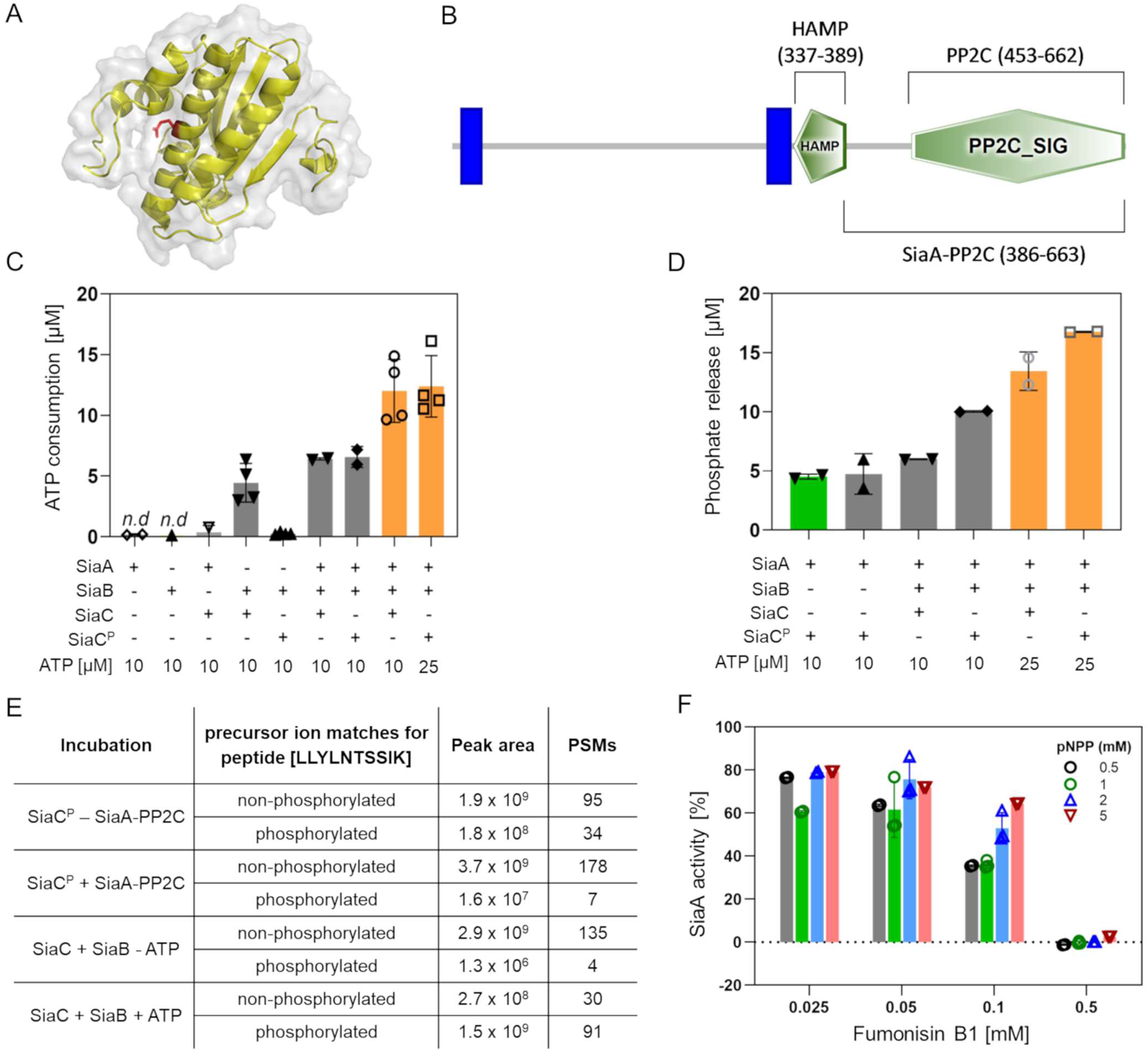
**(A)** Homology model of SiaB (PA0171) predicted using I-Tasser (https://zhanglab.ccmb.med.umich.edu/I-TASSER/) shown as cartoon with 80% transparent surface using the Pymol software (version 2.1.1). The predicted catalytic glutamic acid residue at position 61 is highlighted (*red sticks*). **(B)** Representation of SiaA using the SMART online tool (http://smart.embl-heidelberg.de/). The predicted PP2C_SIG domain, the HAMP domain and the two membrane-spanning regions (*blue bars*) are represented. **(C)** ATP consumption of SiaB measured by the ADP-Glo™ assay kit. Assays were performed for 1 h at 37°C in 20 mM Tris-HCL [pH 7.5], 150 mM NaCl in the absence or the presence of 10 mM ATP. **(D)** Phosphate quantification using a malachite green assay after incubation. Reactions were performed for 1 h at 37°C with 0.5 μM SiaA or SiaB and 5 μM of SiaC or SiaC^P^ in 20 mM Tris-HCL [pH 7.5], 150 mM NaCl buffer containing 20 mM MgCl_2_ or MnCl_2_ (*green bar*), and 10 or 25 μM ATP (*orange bars*). **(E)** Summary of shotgun peptide-mass spectrometry (PMS) results from analysis of SiaC purified from *E. coli* and the Δ*siaA* mutant after incubation with SiaB and SiaA-PP2C, respectively. Incubations were performed for 2 h at 30°C prior to separation by SDS-PAGE for PMS (see Material and Methods) **(F)** Phosphatase activity of SiaA-PP2C in the presence of 0.5-5 mM of pNPP and 0-0.5 mM of fumonisin B1. Assays were performed in 150 mM NaCl, 20 mM Tris-HCl pH 7.5 at 37°C and measured as an increase in absorbance [405 nm]. The initial rate of reaction was calculated and expressed as a percentage compared to the untreated control. Data in **C, D** and **F** are presented as the average from at least two independent enzymatic assays (*bars*; n = 2-4) with the corresponding standard deviation of the mean (*error bars*) and the individual data points (*circles*). Results below the detection limit are indicated (*n.d.*).

Activity measurements using a malachite-green assay (**Fig 4D**) revealed phosphatase activity of SiaA-PP2C in the presence of SiaC^P^ (5 µM; prepared by incubating SiaB/SiaC protein mixture in the presence of MgCl_2_/ATP) and Mg^2+^ (4.74 µM) or Mn^2+^ (4.52 µM) ions. No activity was observed when SiaA was incubated with SiaC. Notably, reactions in which SiaB and SiaC were incubated together with SiaA-PP2C (0.5 µM), the consumption of ATP was increased (6.42 μM and 6.57 μM in the presence of 10 μM ATP and to 12.01 µM and 12.39 µM in the presence of 25 μM of ATP, respectively) compared to SiaB with SiaC in the absence of SiaA-PP2C (**Fig 4C**). Similarly, incubation with SiaA-PP2C, SiaB, and SiaC or SiaC^P^ in the presence of Mg^2+^ ions and 25 μM ATP also indicated much higher phosphatase activities compared to SiaA-PP2C with SiaC^P^ in the absence of SiaB (13.44 µM and 16.76 µM, respectively) (**Fig 4D**). As neither activity correlates with the amount of SiaC/SiaC^P^ added (5 µM), these experiments indicate that SiaA-PP2C and SiaB are both enzymatically active on their respective substrates, catalysing a cycle of SiaC phosphorylation and de-phosphorylation in dependency of ATP availability.

To determine the exact site of phosphorylation, we performed kinase (SiaB + SiaC) and phosphatase (SiaA-PP2C + SiaC^P^) assays and analysed the SiaC/SiaC^P^ protein by PMS after gel purification (**Fig 4E** and **Table S1**). For SiaB with SiaC in the presence of ATP, we observed a 1000-fold increase of the phosphorylated T68-containing peptide (peptide [LLYLN**T**SSIK]) compared conditions in which ATP was omitted from the assay (1.5 × 10^9^ vs. 1.3 × 10^6^ peak area). Incubation of SiaA-PP2C with SiaC^P^ resulted in > 10-fold decrease in the phosphorylated peptide (LLYLN**T**SSIK) compared to the control incubation without SiaA-PP2C added. Hence, the PMS data obtained from these enzyme assays provide direct evidence that T68 of SiaC is a target for phosphatase and kinase activity of SiaA and SiaB, respectively.

### The SiaA activity can be inhibited by fumonisin B1

Given that loss of SiaA function results in poor aggregate and biofilm formation via control of the phosphorylation status of SiaC, SiaA function may represent a novel target for biofilm control through antagonists. Fumonisin B1 is a well-established inhibitor of PP2C type phosphatases [75]. To determine if fumonisin B1 can interfere with SiaA activity, we incubated purified SiaA-PP2C (50 µg/mL) in the presence of different concentrations of the inhibitor as well as varied concentrations of the substrate, pNPP (**Fig 4F**). At increasing concentrations, fumonisin B1 was indeed inhibitory of SiaA phosphatase activity dependent on the concentration of substrate used. In the presence of 0.025 mM fumonisin B1, SiaA-PP2C retained 60.31-79.25% of its activity as compared to the untreated control. At higher fumonisin B1 concentration of 0.1 mM SiaA-PP2C activity was inhibited in a pNPP dependent manner, with 35.48% activity remaining with 0.5 mM pNPP, 39.37% activity with 1 mM pNPP, 52.97% activity with 2 mM pNPP and 64.30% activity retained with 5 mM pNPP used. At 0.5 mM fumonisin, there was complete inhibition of SiaA-PP2C activity across all pNPP concentrations.

### Crystal structure of the PP2C domain of SiaA reveals a homodimer

To further characterise the PP2C domain of SiaA (SiaA-PP2C), the structure of the protein with Mg^2+^ ions at the active site was determined (PDB ID: 6K4E). Native crystals of SiaA-PP2C were obtained in conditions A11 (0.03 M MgCl_2_, 0.03 M CaCl_2_, 0.1 M tris, 0.1 M BICINE, 20 % v/v glycerol, 10 % w/v PEG 40000, pH 8.5) from Morpheus (Molecular Dimensions) with space group of P2_1_2_1_2_1_ and diffracted to a resolution of 2.1 Å. Phases were obtained using single-wavelength anomalous dispersion (SAD) with a single SeMet derivative crystal. Statistics of the data collection, structure determination and refinement are displayed in **Table S2**. The SiaA-PP2C domain is arranged as a tight dimer in the crystal asymmetric unit. The electron density map of SiaA-PP2C allowed unambiguous tracing of residues 407 to 663 for monomer A and 409 to 663 for monomer B. However, no clear electron density was present for residues 521-528 and 578-581 of chain A and residues 522-527 and 577-579 of chain B, indicating a high degree of flexibility in these solvent-exposed loops regions. The SiaA-PP2C monomer adopts the canonical α-β-β-α fold first described for the human PPM1A [76] and subsequently found in several other phosphatases [77,78] (**Fig 5A**). The core structure of the SiaA-PP2C monomer is a β-sandwich composed of two antiparallel β-sheets each comprising five β-strands. The active site is at the apex of the β-sheet structure (**Fig 5A, B** and **D**). This core structure is flanked on the N-terminal side by helices α1-α4 and on the C-terminal end by helices α5-α7, forming a four-layered αββα structure. The two monomers interact extensively with each other through residues projecting from helices α1, α2 and α3, forming a compact homodimer that buries a total solvent accessible area of 1392 Å^2^ (**Fig 5B**). Consistently, in size-exclusion chromatography, instead of monomer (∼33 kDa), SiaA-PP2C eluted as a 66 kDa dimer, suggesting SiaA exists mainly as dimer in solution (**Fig 5C**). The homodimer presents two concave surfaces that are likely to accommodate the incoming substrates (**Fig 5D)**.

**Fig 5:**
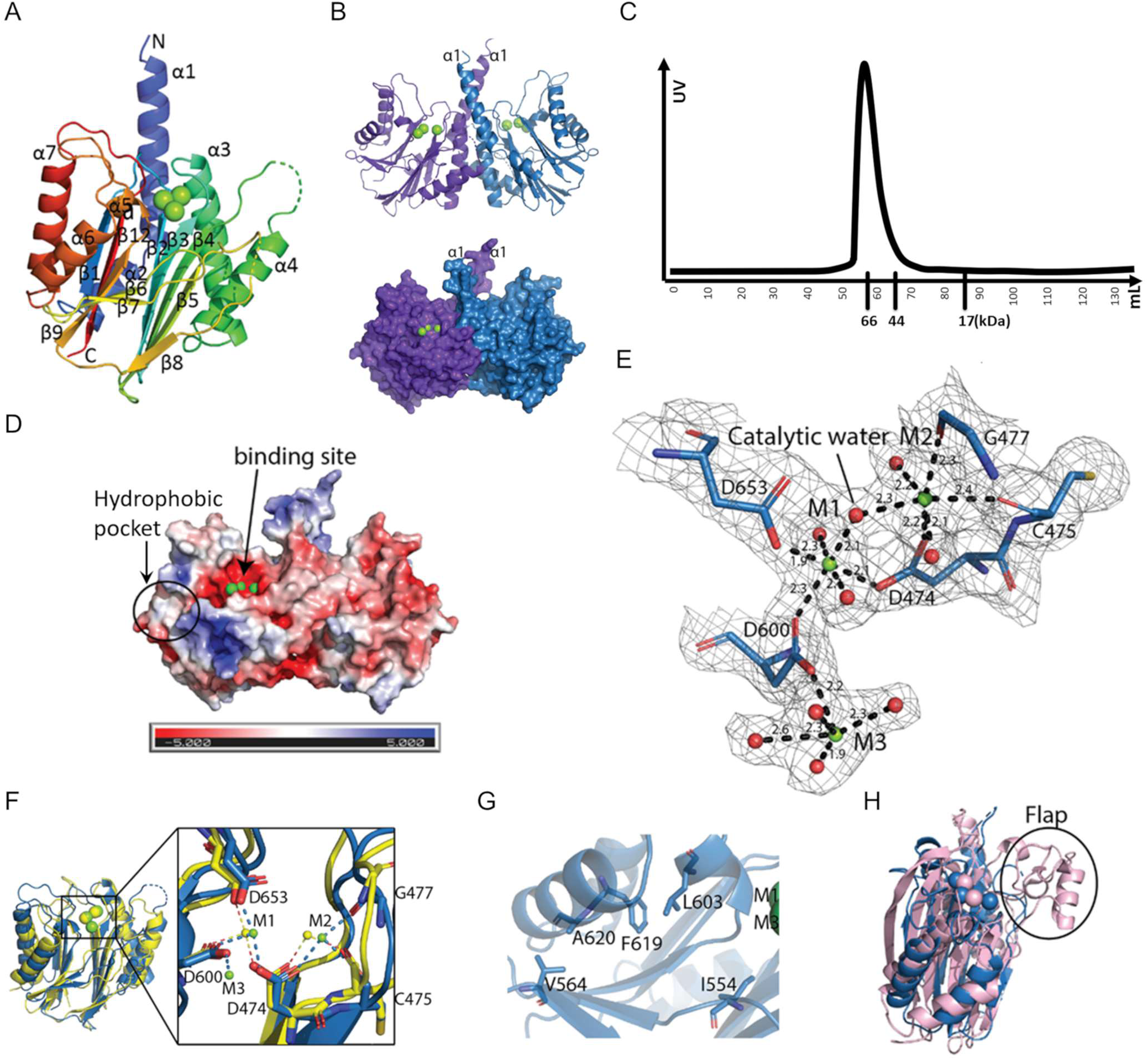
The SiaA-PP2C structure (PDB ID: 6K4E). **(A)** “Rainbow” representation of the SiaA-PP2C monomer. The N-terminus of SiaA-PP2C is coloured in blue and the C-terminus in red, with secondary structures labelled. The three Mg^2+^ ions identified at the active site are represented as green spheres. **(B)** The SiaA-PP2C dimer displayed in cartoon (top) or surface (bottom). Monomer A is in purple and B in blue. **(C)** Size-exclusion chromatography of SiaA-PP2C. Elution volumes of markers are labelled below X-axis. **(D)** APBS surface electrostatic potential of SiaA-PP2C homodimer in the same orientation as in panel B. **(E)** Close-up view of the active site of the PP2C-like phosphatase domain of SiaA-PP2C with conserved acidic residues in the active site shown as sticks and labelled. An electron density map with Fourier coefficients 2F_O_-F_C_ is overlaid at a level of 1σ above the mean. The Mg^2+^ ions are shown as green spheres and the oxygen atoms of the coordinating water molecules are displayed as red spheres. M1 and M2 are both coordinated by oxygen atoms from three residues and three water molecules in an octahedral manner. Distances between the metal ions and the coordinating atoms are indicated. **(F)** Overlay of SiaA_PP2C monomer B (*blue*) and the phosphatase domain of Rv1364C (*yellow*). Mg^2+^ ions M1 and M2 of SiaA-PP2C shown as green spheres overlap with Mn^2+^ ions of Rv1643C (*yellow spheres*). The α1 and α2 helices of SiaA-PP2C are not displayed for clarity. **(inset)** Magnified view of the active site. Metal coordinating residues of SiaA-PP2C are labelled. **(G)** Residues of the hydrophobic pocket shown as sticks. **(H)** Superposition of SiaA monomer B (*blue*) and human PPM1A (*light pink*; PDB code 6B67-A). The Flap sub-domain of human PPM1A (absent in SiaA-PP2C) is circled.

An automated search of 3D structures similar to SiaA-PP2C, using the Dali server (ekhidna2.biocenter.helsinki.fi/dali), returned PP2C-type phosphatases as the top matches, the best of which was Rv1364C (PDB code: 3KE6, chain A) from *Mycobacterium tuberculosis* with a Z score of 23.6 and a RMSD of 2.2 Å for 210 superimposed α-carbon atoms. Despite a very low overall amino-acid sequence identity (17%), these structures share a conserved fold and several strictly conserved active site residues with SiaA-PP2C: D457, D474, G477, D600, G601 and D653 (**Fig 5E** and **Fig S2**). A structural comparison of the active site of SiaA-PP2C with the active site of the phosphatase domain of Rv1364C is shown in **Fig 5F**. The two Mg^2+^ ions labelled as M1 and M2 directly involved in the catalytic mechanism are coordinated by oxygen atoms from three evolutionary conserved aspartate residues and occupy equivalent positions in SiaA-PP2C compared to the two Mn^2+^ ions found in the Rv1364C active site. Mg^2+^ ion was placed here because 0.03 M MgCl_2_ was present in crystallization buffer, and hence it does not indicate SiaA-PP2C’s preference for metal ions. Mn^2+^ and Mg^2+^ has both been reported as ligand of PP2C, and it is possible that Mn^2+^ and Mg^2+^ both function as co-factor of SiaA-PP2C. Metal-coordinating residues are all conserved except C475 of SiaA-PP2C, which interacts with metal ion M2 through its carbonyl oxygen (**Fig 5E** and **F**).

Compared to Rv1364C, which only has two bound metal ions, a third Mg^2+^ ion (M3) was found in SiaA-PP2C coordinated by D600 and with an incomplete octahedral coordination shell. D604 and D534 assist in stabilization of M3 through formation of hydrogen bond with M3-coordinating water molecules. Given the structural similarity with PP2C [76], an S_N_2 mechanism is the most plausible whereby the water molecule bridging M1 and M2 (**Fig 5E**) performs a nucleophilic attack of the phosphorus atom from the phosphate bound to the threonine target residue. However, M3 could also play a direct role in the catalytic activity, as was recently proposed for the human PPM1A protein, a negative regulator of cellular stress response pathways [79,80]. Further study is needed to investigate whether a third metal ion is present in all PP2C. We note that near the entrance of the active site, a hydrophobic pocket lined with residues I554, V564, L603, F619 and A620 constitutes a potential site for the design of inhibitors (**Fig 5D** and **G**). Another distinct feature of SiaA-PP2C is a flexible loop located between β7 and β8, whereas a Flap subdomain, which has been reported to aid in substrate specificity, is present at the equivalent position in human PPM1A and many other PP2C-type phosphatases [80] (**Fig 5H**). Therefore, this shorter yet flexible loop might contribute to the substrate specificity of SiaA-PP2C.

### Crystal structure of SiaC reveals similarity with anti-sigma factor antagonists

To gain insights about SiaC, its crystal structure was determined in this study. Native crystals of SiaC were grown in C6 condition of the JCSG+ kit from Molecular Dimensions (0.1 M Phosphate/citrate, pH 4.2, 40% (v/v) PEG 300), with space group I222 and diffracted to a maximum resolution of 1.7 Å. The structure (PDB ID: 6K4F) was determined by SAD using SeMet derivative crystals (**Table S2**).

Clear electron density maps allowed complete tracing except for residues 101-103, which are located in the flexible loop between the α3 helix and β5 strands. The SiaC protein comprises six β-strands arranged in a mixed parallel/antiparallel fashion and three α-helices (**Fig 6A**). A hydrophobic pocket next to T68 is formed by residues L61 and L63 projecting from β4, W95 from β5, I71 and M74 from α2, L107 and F111 from α3 while L66 belongs to the β4-β5 loop (**Fig 6B** and **C**). Dali server was used to search for protein structures that are most similar to SiaC crystal structure, and an anti-sigma factor antagonist from *Mycobacterium paratuberculosis* (PDB ID: 4QTP; gene ID MAP_0380; **Fig 6D**) was identified as the closest homologue (Z score of 9.0, an amino-acid sequence identity of 10% and a RSMD of 2.8 Å over the □-carbon atoms of 117 residues). Despite sharing very low sequence identity, a closer examination of both structures reveals a conserved overall topology with several unique features, especially at β3, β4 strands and α3 helix that are longer in 4QTP compared to SiaC. These variations are probably related to differences in the ability to establish protein-protein interactions. Nonetheless, their target phosphorylation sites, T68 for SiaC and S58 for 4QTP, are both located at the N-terminus of helix α2.

**Fig 6:**
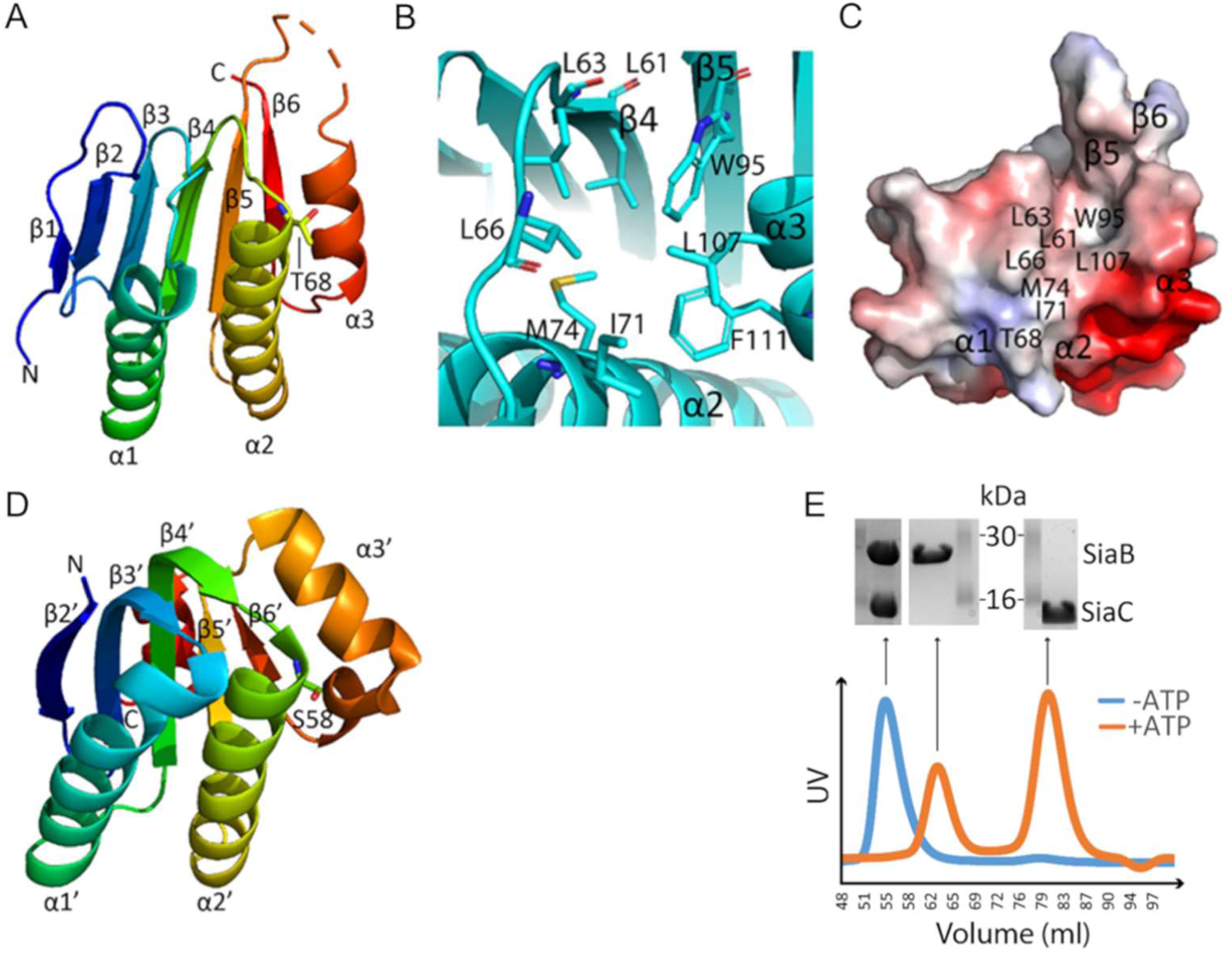
**(A)** “Rainbow” representation of SiaC (PDB ID: 6K4F). The phosphorylation site T68 is shown as sticks. **(B)** Hydrophobic pocket of SiaC next to T68. **(C)** Surface electrostatic potential of SiaC with the hydrophobic pocket labelled. The structure is displayed in the same orientation as in panel B**. (D)** “Rainbow” representation of 4QTP. The phosphorylation site S58 is shown as sticks. **(E)** Size-exclusion chromatography profiles derived by UV-Absorption for SiaB/SiaC mixtures (1:1 molar ratio) in the presence (*orange line*) or absence (*blue line*) of 10 mM ATP and 20 mM MgCl_2_. The SDS-PAGE analysis of the proteins causing the peaks in the size exclusion chromatography are shown above the profiles.

It is worth mentioning that our SiaC crystal structure not only agrees with the SiaC structure published by Chen et al. [73], but also provides more details at 1.7Å resolution. A 2SiaB:2SiaC complex was suggested by size-exclusion chromatography when SiaB and SiaC were mixed in equal molar ratio and incubated at room temperature for 1 h (**Fig 6E**). Instead of two individual peaks (one consists of SiaB and one consists of SiaC), a single peak containing the complex was observed. This 2SiaB:2SiaC complex model is consistent with the recently reported crystal structure of the SiaB/SiaC complex [73]. When the SiaB/SiaC mixture was supplemented with 10 mM ATP and 20 mM MgCl_2_ prior to size-exclusion chromatography, the complex disappeared and SiaB and SiaC^p^ eluted as two individual peaks. The phosphorylation state of SiaC^P^ was subsequently confirmed by LC-MS. As such, our data demonstrate that SiaB binds tightly to SiaC but quickly dissociation from SiaC^P^ after phosphorylation.

### Molecular Docking and MD simulation of SiaC^P^ with the phosphatase domain of SiaA

As shown above, the SiaB/SiaC complex quickly dissociates upon phosphorylation. Since SiaA-PP2C targets SiaC^P^, we wanted to study the interaction between SiaA and SiaC during the dephosphorylation reaction. Therefore, we performed a MD simulation on SiaA binding to SiaC and to SiaC^P^. The SiaA/SiaC^P^ complexes were predicted by molecular docking of SiaA-PP2C dimer to SiaC and to SiaC^P^ using ZDOCK, followed by 60 ns of MD simulation (**Fig 7A** and **B**). These simulations suggest that the phosphorylated pT68 is required for productive binding to the catalytic site of SiaA-PP2C, where it remains efficiently bound via the catalytic Mg^2+^ ion M3 and the bridging water molecules surrounding it (**Fig 7C-E**). This result is consistent with the experimental observation that only pT68 in SiaC^P^ constitutes the substrate for SiaA-PP2C activity. In contrast, the non-phosphorylated T68 dissociated from the catalytic site in less than 15 ns of simulation, suggesting that the interaction of SiaC with SiaA is weak upon dephosphorylation. Based on the simulated model, several key interactions that could contribute to substrate specificity of SiaA-PP2C were identified. First, R652 stabilizes the positively charged phosphate group of pT68 through the formation of a salt bridge (**Fig 7F**). In addition, the flexible loop between α5 and α6 of SiaA inserts into a shallow pocket located between α2, α3, β4, β5 and β6 of SiaC^P^ and establishes hydrophobic contacts with L61, L66, I71, M74, M75, L78, W95, L107 and F111 through F612 (**Fig 7G**). Notably, the salt bridge identified between R618 and E609 of SiaA is likely to restrain the orientation of the flexible loop between α5 and α6, which could favour interaction with SiaC^P^. Notably, F612 was recently found to be deleted in the SiaA protein from a non-aggregative mutant (Δ*siaA*; G611F612; SiaA-PP2C*)[70].

**Fig 7:**
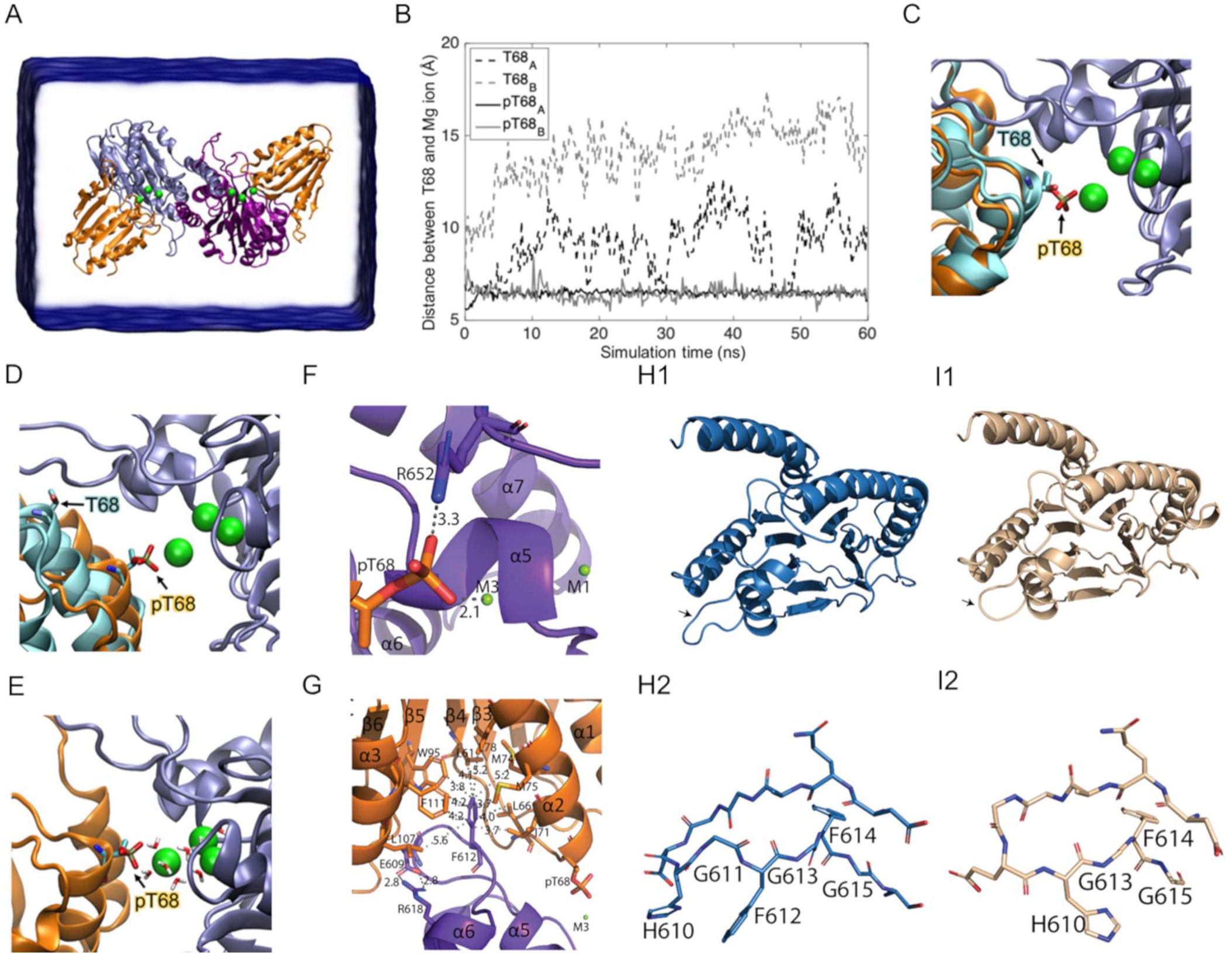
Binding of SiaA-PP2C to phosphorylated SiaC. **(A)** The simulated system using ZDOCK contains a SiaA-PP2C dimer (*blue/purple ribbon*) with SiaC (*cyan ribbon*) or SiaC^P^ (*orange ribbon*) docked to each SiaA-PP2C monomer. **(B)** Distance between Cα of T68/pT68 and Mg^2+^ ion (M3). Phosphorylated T68 of both SiaC^P^ (*black and grey solid lines*) remains coordinated with the Mg^2+^ ion (*green sphere*). Unphosphorylated T68 (*black and grey dashed lines*) dissociates from the Mg^2+^ ion in < 15 ns. The positions of T68 and Mg^2+^ ions before **(C)** and after the simulation **(D)** are displayed in the same color code as in panel A. **(E)** The pT68 of SiaC^P^ is interacting with the bridging water molecules for the entire 60 ns of simulation. **(F-G)** Cartoon representations of specific parts of the SiaA-PP2C/SiaC^P^ complex after 60 ns of simulation (**Supplementary zip-file**). SiaC^P^ is coloured in orange; SiaA-PP2C dimer is coloured in purple and blue **(F)** Interaction of pT68 with R652 and M3. R652 might be important for stabilisation of the complex through the formation of a salt bridge. pT68 remains in close proximity with M3 (2.1 Å). **(G)** F612 interacts with the hydrophobic pocket of SiaC^P^. Salt bridge formation between R618 and E609 in SiaA-PP2C might restrain the orientation of the flexible loop between α5 and α6 to accommodate its interaction with the hydrophobic pocket of SiaC^P^. Cartoon representation of SiaA-PP2C **(H1)** and a homology model of the mutant allele SiaA-PP2C* **(I1)** from the Δ*siaA* mutant (**Supplementary zip-file**), in which G611 and F612 at the flexible loop between α5 and α6 are deleted (*black arrows*). Stick representation of the flexible loop of SiaA-PP2C **(H2)** and SiaA-PP2C* **(I2)**.

A homology model of SiaA-PP2C*, (**Supplementary zip-file**) using SWISS-MODEL (https://swissmodel.expasy.org/) with the SiaA-PP2C crystal structure as template (GMQE =0.98, QMEAN = −0.54) confirmed that the deleted amino acids G611F612 most likely impacts the orientation of the flexible loop between α5 and α6, and confirmed that F612 seems to be important for the interaction with the hydrophobic pocket of SiaC^P^ **(Fig 7H1-I2)**. As such, a stable SiaC^P^-SiaA-PP2C* complex seems unlikely to form, which would explain the non-aggregative phenotype of the Δ*siaA* mutant strain due to an impaired enzymatic activity towards SiaC^P^.

## Discussion

The SiaA and/or SiaD proteins are essential for the formation of macroscopic aggregates during growth in the presence of the toxic surfactant SDS or the ROS-generating biocide tellurite in a c-di-GMP manner [7,70,72]. This active and energy requiring response may represent an adaptive survival strategy as the corresponding aggregates were found to protect cells of the same, as well as other, species in mono- and mixed culture experiments [11,71,81]. Here, we show that the SiaABCD proteins play a key role in cellular aggregation and dispersal of *P. aeruginosa* in response to carbon availability in the environment. This is intriguing in the context of lung colonisation by *P. aeruginosa* given the dynamic changes in the sources and supply of carbon substrates available in the lung, including but not limited to, glucose [82], short-chain fatty acids [83,84], ethanol and 2,3-butanediol [85–87]. Notably, ethanol and 2,3-butanediol have recently been reported to influence microbial colonisation of epithelial cells or the persistence of microbes during infection [13,88,89]. It is thus possible that the SiaABCD signalling pathway is not only involved in the regulation of cellular aggregation but also has an impact on the virulence of *P. aeruginosa*. This hypothesis is supported by several observations:

i. A diverse set of regulatory systems involved in biofilm formation and/or virulence are interconnected with the SiaABCD pathway. These include the transcriptional regulators AmrZ and FleQ or the posttranscriptional regulatory systems RsmA/*rsmY*/*rsmZ*, RsmN and CRC [72,90–94].
ii. SiaD modulates the community structure and competitiveness of *P. aeruginosa* cells within dual-species biofilms with cells of *Staphylococcus aureus* and impacts pyoverdine and pyocyanin production [7,71,95].
iii. SiaD was identified as one of the DGCs capable of initiating the “Touch-Seed-and-Go” virulence program by activating the c-di-GMP receptor FimW [96]. This activation leads to the attachment of cells and the subsequent induction of virulence traits through asymmetric cell division, finally promoting efficient tissue colonization, localised host damage and fast dissemination of the infection.

With our study, we demonstrated that the protein phosphatase SiaA and its kinase counterpart SiaB target the protein SiaC at threonine 68 (T68) and provide evidence that the SiaABC proteins make use of a partner switch mechanism to balance cellular aggregation in response to carbon availability. A partner switch system (PSS) typically involves a PP2C phosphatase (SiaA), an anti-sigma factor with kinase activity (SiaB) and anti-sigma factor antagonist representing the target for phosphorylation (SiaC). In line with such a mechanism, the phosphorylation status of the proposed anti-sigma factor antagonist SiaC is crucial for the switch to occur. In addition to the sequestration of an alternative sigma factor, partner switch systems have recently been found to regulate important physiological traits also by affecting enzymatic activities involved in cellular signalling cascades through direct protein-protein interactions.

In *P. aeruginosa*, the HptB-HsbR-HsbA system was shown to promote DGC activity of HsbD following its interaction with the phosphorylatable anti-sigma factor antagonist HsbA to regulate biofilm formation and swimming motility [97]. More recently, a PSS was shown to be involved in mixed-linkage ß-glucan synthesis in *S. meliloti* through the induction of c-di-GMP levels [98]. In this system, the Ser/Thr-specific phosphatase/kinase couple BgrU/BgrW controls the phosphorylation status of the BgrV protein and hence controls the activity of the DGC BgrR. Finally, in *Moorella thermoacetica* the RsbT-like switch kinase MtT was found to bind to and attenuate the activity of the DGC MtG (PDB: 3ZT9) after phosphorylation of the RsbS-like stressosome scaffold protein Mts (Mts-P [PDB: 3ZTB]). Notably, SiaD activity exhibits the same phenotype as loss of SiaC function (this study), the *siaABCD* genes are co-regulated in an operon [71] and SiaD overexpression was found to be incapable of complementing the non-aggregative phenotype of the Δ*siaA* mutant strain whereas other DGCs could [70]. Based on these facts, one can speculate that the SiaABC partner switch functions to regulate SiaD mediated c-di-GMP biosynthesis through the phosphorylation status of SiaC. In one such scenario, the activation of SiaA would shift the equilibrium of SiaC/SiaC^P^ towards the unphosphorylated form of SiaC, which subsequently interferes with SiaD activity, leading to increased c-di-GMP production and subsequently biofilm formation. Such a scenario would explain the strong aggregative phenotype of the SiaB mutant observed in this and previous studies [99]. As a negative feedback response, SiaC^P^ has an increased binding affinity towards SiaA, which would lead to a negative impact on SiaD activity through the dephosphorylating of T68 and increased binding to SiaB.

During the publication of our preprint [100] and the per-review process for this manuscript, Chen et al. [73] published a similar data-set that showed direct interaction of SiaC with SiaB and SiaD based on two-hybrid data and co-crystallization of the SiaB-SiaC complex as well as activation of SiaD through SiaC binding. As such, that study is complementary to our data and strongly supports a model that the SiaABC partner switch integrates environmental information, such as carbon availability and surfactant stress, to control biofilm formation through the modulation the activity of SiaD through the phosphorylation status of SiaC.

Given their importance in regulating aggregation and biofilm formation, the PP2C domain of SiaA and the putative anti-sigma factor antagonist SiaC may represent novel targets for the development of biofilm interference drugs. The PP2C-type phosphatase SiaA, while conserved in prokaryotes and eukaryotes, is structurally unique. A Flap sub-domain, suggested of being involved in substrate interaction and specificity, has been reported in human PPM1A and many other PP2C-type phosphatases [80]. SiaA-PP2C lacks the Flap sub-domain and instead has a long loop between β7 and β8 that does not seem to interact with SiaC^P^ according to MD simulations. However, the flexible loop between α5 and α6 might aid in substrate specificity of SiaA by interacting with the hydrophobic pocket of SiaC near the phosphorylation site. Although further studies are needed to confirm this finding, the fact that the phosphatase domain of the mutant allele SiaA-PP2C* has a deletion in G611F612, of which the latter residue seems to be crucial for the interaction with SiaC, is supportive of this hypothesis. Thus, this flexible loop in SiaA could be exploited for drug design to interfere with SiaA/SiaC^P^ interaction and hence, block downstream induction of biofilm formation through the inhibition of SiaC^P^ dephosphorylation. In addition, we demonstrated that the phosphatase inhibitor fumonisin B1 [75] was capable of completely inhibiting the activity of SiaA *in vitro*. Fumonisin B1 is also known to inhibit ceramide synthase [101], and it was proposed to work by establishing electrostatic interaction between negatively charged tricarboxylic groups and positively charged active site. Here we speculate that fumonisin B1 might inhibit SiaA through a similar mechanism, namely by electrostatic interactions between the negatively charged tricarboxylic groups of fumonisin B1 and the positively charged metal ions of SiaA, thus blocking the active site. In addition to SiaA inhibition, SiaC also represents an exciting target for interference, as it is structurally most similar to bacterial anti-sigma factor antagonists with little homology with human proteins. The hydrophobic pocket next to the phosphorylation site of SiaC is of particular interest, as it seems to be involved in the stability of the SiaA/SiaC^P^ interaction. In addition, a large patch of negatively charged surface is present at α2 and α3 near the hydrophobic pocket **(Fig 6C)**, and could thus be useful for drug design as potential anchor points.

In summary, the presented work provides a deeper understanding of the SiaABCD mediated regulation by expanding the fundamental genetic and biochemical mechanisms that control cellular aggregation in response to carbon availability. Together with the insights into the catalytic and regulatory mechanisms of the protein-phosphatase family-2C (PP2C) domain of SiaA and of the SiaC protein gained by our structural analysis, the present study might facilitate the development of future anti-biofilms drugs based on a signal interference strategy, which are desperately needed to combat the rising crisis due to antibiotic resistance.

## Materials and Methods

### Strains and growth conditions

Bacterial strains (**Table S3**) were maintained on Lysogeny Broth (LB) (Bertani 1951) solid medium (1.5% agar w/v) and routinely cultured in 10 mL LB medium in 50 mL Falcon tubes (Greiner Bio-one) or M9 medium (47.6 mM Na_2_HPO_4_, 22 mM KH_2_PO_4_, 8.6 mM NaCl, 18.6 mM NH_4_Cl, 2 mM MgSO_4_, 0.1 mM CaCl_2_, 0.03 mM FeCl_2_) with shaking at 200 rpm and incubation at 37°C or 30°C.

### Strain construction

To create the *siaB* and *siaC* mutant strains, the Tet^r^ IS*phoA*/*hah* transposon insertion located in the *siaB* and *siaC* genes was transferred into the wild type strain by E79tv2 phage transduction [24], using the PW1292 and PW1290 strains from the University of Washington Genome Centre as the source of the transposon [102] (**Table S3**). Following transduction, Tet^r^ resistant strains containing IS*phoA*/*hah* insertions were cured of the antibiotic resistance marker by Cre/*loxP* recombination mediated excision of the tetracycline resistance cassette. For expression of the Cre recombinase, the pCre1 plasmid was introduced into the *P. aeruginosa* mutant strain harbouring the IS*phoA*/*hah* insertion by conjugation using biparental mating with S17-λpir. An overnight culture of the *P. aeruginosa* recipient strain was grown in 10 mL LB at 42°C with shaking at 50 rpm, while the *E. coli* donor strain was grown in 10 mL LB at 37°C with shaking at 150 rpm. Optical density measurements were used to provide 2 × 10^9^ cells of the recipient strain and 1 × 10^9^ cells donor strain, where an OD_600_ = 1 is approximately 1 × 10^9^ cells/mL. Cells were washed twice in 1 mL pre-warmed LB with centrifugation for 30 sec at 10,000 × *g* at room temperature (Heraeus instruments; biofuge pico). Finally, the cells were combined in 100 µL LB pre-warmed to 37°C, spotted on to a hydrophilic mixed cellulose 0.45 µm sterile filter (Pall Corporation) on a LB plate pre-warmed to 37°C and incubated at 37°C for 8 h for transfer. Following incubation, the filter was transferred to a 50 mL Falcon tube and the bacteria were resuspended in 1 mL 0.9% NaCl (w/v) by vortexing. Serial dilutions were performed in M9 medium and subsequently plated on Pseudomonas Isolation Agar (PIA, Difco) plates to counter-select against *E. coli* cells and to determine the colony forming units (CFU/mL). Based on the CFU results, appropriate dilutions were selected to ensure that 50-100 colonies from the initial transduction were spread on to the PIA agar plates. After growth for 14-16 h at 37°C, single colonies of the exconjugants were patched on to both selective and non-selective agar plates to test for tetracycline sensitivity. The correct site of insertion and loss of the tetracycline resistant cassette was confirmed by PCR and sequencing.

### Plasmid construction

Cloning of the C-terminal part of SiaA (SiaA-PP2C), SiaB, and SiaC were carried out using ligation-independent cloning methods as described in Savitsky et al. [103]. The gene regions encoding SiaA-PP2C (amino acids 386-663; Genbank ID: NP_248862), SiaB (Genbank-ID: 248862), SiaC (Genbank-ID: NP_248860) were amplified via PCR of genomic DNA of the wildtype strain with appropriate primers (**Table S4**). Amplified sequences were treated with T4 DNA polymerase and dCTP. The vector pNIC28-Bsa4 was digested with *Bsa*I whereas the vector pNIC-CTHF was digested with *Bfu*AI. The vectors were then purified and subsequently treated with T4 DNA polymerase and dGTP. Treated vectors and inserts were mixed and annealed for 10 min at room temperature before being used for transformation of Mach1 chemically competent *E. coli* cells leading to the expression vectors pNIC[SiaA], pNIC[SiaB] and pNIC[SiaC], respectively. For protein expression, the construct was retransformed into *E. coli* BL21(DE3) Rosetta T1R competent cells. Small scale analytical expression screens were then carried out to verify expression of the protein and its likelihood for success in large scale purification. Transformed *E. coli* BL21(DE3) Rosetta T1R were grown at 37°C in 1 mL of Terrific Broth (TB) (1.2% w/v tryptone, 2.4% w/v yeast extract, 0.5% v/v glycerol, 89 mM phosphate buffer) to an OD_600_ of ∼ 2 −3. The temperature was subsequently reduced to 18°C and protein production was induced with 0.5 mM Isopropyl-β-D-thiogalactoside (IPTG, Sigma Aldrich) overnight. The cells were then lysed and subjected to clarification and microscale immobilized metal-ion affinity chromatography (IMAC) to assess the levels of soluble and purifiable target protein from each clone.

For construction pJEM[SiaC], the His6-TEV-*siaC* construct from pNIC[SiaC] was amplified using Q5 Hot-Start DNA polymerase (New England BioLabs) with the appropriate primer pair (**Table S4**). The primers had 25 bp of homology to the insertion sites in plasmid pJeM1 on each site (marked as bold text). The eGFP gene in plasmid pJeM1 was excised using *Nde*I and *Hind*III and replaced with the purified PCR products *via* the one-step isothermal assembly. The constructs were subsequently transformed into *E. coli* DH5α (Invitrogen) and the correctness of the cloned genes was verified by sequencing (GATC Biotech).

### Phenotypic characterisation in 12 well plates

For phenotypic characterisation of cultures during growth on 3.5 mM SDS and 22.2 mM glucose, cell culture treated, 12 well microtiter plates with 2 mL M9 medium were prepared and inoculated with washed cells from an overnight culture grown in LB medium to an OD_600_ = 0.01. Following incubation at 30°C with shaking at 200 rpm for 18 h the plates were imaged at 1200 dpi using an Umax Powerlook 1000 flatbed scanner with V4.71 software in a dark room. Images were normalised using Adobe Photoshop CS5 Software by using the ‘match colour’ image adjustment function, with all images normalised to the first image scanned using this experimental format. Experiments were performed as at least 6 independent replicates and representative images are shown.

### Analytics of supernatants

For glucose quantification, the liquid cultures were filtered through a 0.22 µM PES filter (PN 4612, Pall) and quantified using the GO assay kit (GAGO20, Sigma Aldrich). Briefly, one volume of sample or standard solution (20 µL) was mixed with two volumes of assay reagent (40 µL) in a 96 well plate and incubated statically at 37°C for 30 min. Subsequently, the reaction was stopped by the addition of 12 N H_2_SO_4_ (40 µL) to the reaction mixture. Glucose concentrations were quantified at 540 nm using a microplate reader (Infinite 200 pro, Tecan). Zero-80 µg/mL glucose standard solutions were used for the calibration curve.

Concentrations of succinate, ethanol and 2,3-butanediol were quantified by a HPLC method using an Aminex HPX-87H 300 × 7.8 mm ion-exchange column (BioRad, Munich, Germany) heated to 60°C. The eluent was 5 mM H_2_SO_4_, which was delivered to the column by a LC-10ATvp pump (Shimadzu, Munich, Germany) at a flow rate of 0.6 mL/min. The eluent was continuously degassed with a DGU-20A3R degassing unit (Shimadzu, Munich, Germany). Samples were injected using 10 µL with a 234 autosampler (Gilson, Limburg-Offheim, Germany). Resolved compounds were analysed with a refractive index detector (RID-10A, Shimadzu, Munich, Germany) and the data processed using the Shimadzu Lab solutions software version 5.81. Concentrations were finally calculated from calibration curves of the corresponding metabolite of interest.

### Biofilm quantification using crystal violet staining in 24 well plates

A microtitre based crystal violet (CV) staining assay was used to quantify biofilm formation during growth with various media. Cell culture treated and non-treated 24 well microtiter plates with 800 µL M9 medium were prepared and inoculated with washed cells from an overnight culture grown in LB medium to an OD_600_ = 0.01. M9 medium was supplemented with 22.2 mM glucose, 20 mM succinate, 40 mM ethanol or 20 mM 2,3-butanediol and incubated at 30°C with shaking at 200 rpm. Following incubation, liquid cultures were removed, pooled and used for quantification of the carbon source and optical density measurements. The wells of the microtitre plates were then stained with 900 µL CV solution (0.1% [w/v] stock in water) for 10 min without shaking. After incubation, the liquid was discarded and the wells of the microtitre plates were washed with 1 mL Tris-HCl (10 mM; pH 7) containing 0.9 % NaCl (w/v) for 10 min, once without and once with shaking at 200 rpm. Subsequently, the plates were air dried and the remaining CV was solubilised by incubation with 1 mL of pure ethanol for 10 min with shaking at 200 rpm. Quantification of CV was quantified at 595 nm in a microtiter plate reader.

### Scanning electron microscopy (SEM)

For SEM, overnight cultures of *P. aeruginosa* strains were prepared in 10 mL LB in 50 mL Falcon tubes and incubated for 16 h at 30°C with shaking at 200 rpm. Following incubation, cultures were harvested by centrifugation for 2 min at 16,060 × *g* at room temperature and washed twice in 1 mL of fresh M9 minimal medium. These cells were used to inoculate 50 mL of medium supplemented with 22.2 mM glucose with an OD_600_ = 0.05 in a 250 mL Erlenmeyer flask and incubated for 5.5 h at 30°C with shaking at 200 rpm. Experiments were performed as three independent replicates, which were subsequently pooled together. Pooled samples were pipetted onto poly-L-lysine coated coverslips on the bottom of the wells of a 12 well plate, using enough culture to create a convex droplet without flooding past the edges of the coverslip. After incubation for 5 min at room temperature, the coverslips were gently flooded with 2 mL M9 minimal medium, which was then removed by a pipette. A fixative solution (glutaraldehyde 2.5%, paraformaldehyde 2%, in M9 buffer) was immediately added to the samples and left overnight at 4°C. The relatively large size of particles formed by the Δ*siaB* mutant exceeded the binding capacity of the poly-L-lysine coverslips, so an alternative fixation process was employed for this strain. After pooling of the three biological replicates, 3 mL of culture was added to a well of a 12 well plate and aggregates were allowed to settle to the bottom of the well by gravity. Once the aggregates had sedimented, the growth medium was removed with a pipette, the cells were washed once in 2 mL M9 medium by this technique and the fixative solution (glutaraldehyde 2.5%, paraformaldehyde 2%, in M9 buffer) was added to the remaining aggregates. Following overnight incubation in fixative solution, all samples were washed 3 times in 2 mL of M9 medium with 5 min incubations at room temperature for each wash. Finally, fixed aggregates of the Δ*siaB* mutant were loaded into a microporous chamber with a 120-200 µm pore size (ProSciTech). Samples of both coverslips and the microporous chamber were then dehydrated by a series of washes with increasing concentrations of ethanol (30%, 50%, 70%, 80%, 90% and 95%), with 5 min incubations at room temperature for each step. Samples were then washed 3 times in 100% ethanol with 5 min incubations at room temperature for each step to complete dehydration. Following dehydration, samples were then dried at the critical point with CO_2_ as the transitional medium (Tousimis, Autosamdri-815). Coverslips were subsequently transferred on to 12.5 mm SEM stubs using carbon impregnated double-sided adhesive tape and left overnight in a 25°C oven for de-gassing of the tape. Microaggregates of the Δ*siaB* mutant were removed from the chamber and placed directly on to the carbon impregnated double-sided adhesive tape on a 12.5 mm SEM stub and left overnight in a 25°C oven for de-gassing of the tape. Samples were then sputter coated with gold for 2 min 30 sec at 40 mA (Emitech, K550X). Conductive samples were then analysed by scanning electron microscopy using a spot size of 3, voltage at 15 kV and a working distance of 10 mm (FEI Quanta 200 ESEM). Multiple fields of view were imaged randomly for each sample and representative images are shown.

### Particle size experiments using laser diffraction analysis (LDA)

Strains were inoculated in 20 mL LB medium (244620, BD Difco) in a 100 mL Erlenmeyer flask and incubated at 30°C with 200 rpm shaking. Overnight cultures were subsequently aliquoted to 50 mL falcon tubes and centrifuged at 9500 × *g* for 5 min. The supernatant was discarded and the pellet was resuspended in 10 mL of M9 medium containing 22 mM glucose. The washing step was repeated once with the pellet resuspended in 2-5 mL of M9 medium. Finally, the cell suspensions were passed five times through a 25 gauge syringe needle to disrupt cell aggregates. The optical density of the suspension was adjusted to OD_600_ = 2 and 1 mL was added to 20 mL of M9 medium supplemented with glucose in an 100 mL Erlenmeyer flask at a final OD_600_ = 0.1. The cultures were subsequently incubated at 30°C with shaking at 200 rpm. Samples were collected at 3 and 6 h for OD_600_ measurement and quantification of glucose concentration. After 6 h of incubation, when glucose was not yet exhausted from the medium, the cultures were analysed with a particle size analyser (SALD 3101, Shimadzu) using a pump speed of 4.0, and the accompanying software was used for analysis of the data (WingSALDII version 3.0.0). Particle sizes of between 0.5-3000 µM were selected for analysis. Calculations were based on water as the dispersing agent with a refractive index of 1.70-020i. Data are presented as the cumulative percentage of the biovolume of the particles within a given size range (0.5-10 μM, 10-200 μM, 200-3000 μM) compared to all particle in the sample. Two biological replicates were analysed and from each sample, the particle size distribution was quantified with three subsequent measurements. Data are presented as the mean value of technical triplicates from a single experiment with the error representing the technical variation.

### Protein production and purification from Escherichia coli cells

The production of the C-terminal part of SiaA (SiaA-PP2C), SiaB and SiaC was performed in *E. coli* BL21(DE3) Rosetta T1R cells harbouring the inducible protein production plasmids pNIC28-Bsa4 and pNIC-CTHF and was performed in a LEX system (Harbinger Biotech). Using glycerol stocks, inoculation cultures were started in 20-40 mL TB medium supplemented with appropriate antibiotics. The cultures were incubated at 37°C with 200 rpm shaking overnight. The following morning, bottles of 0.75-1.5 L TB supplemented with appropriate antibiotics and 100-200 μL of antifoam 204 (Sigma-Aldrich) were inoculated with the overnight cultures. The cultures were incubated at 37°C in the LEX system with aeration and agitation through the bubbling of filtered air through the cultures. When the OD_600_ reached ∼2, the temperature was reduced to 18°C and the cultures were induced after 30 to 60 min with 0.5 mM IPTG. Protein expression was allowed to continue overnight. The following morning, cells were harvested by centrifugation at 15°C and 4000 × *g* for 10 min. The supernatants were discarded and the cells were suspended in protein buffer A (20 mM HEPES pH 7.5, 300 mM NaCl, 5% v/v glycerol, 0.5 mM TCEP) supplemented with 25 μL of protease inhibitors cocktail III (539134, Calbiochem, Merck).

For protein purification, the cell suspensions were sonicated (Sonics Vibra-cell) at 70% amplitude and a 3 sec on/off cycle for 3min, on ice. The lysate was clarified by centrifugation at 47000 × *g* at 4°C for 25 min. The supernatants were filtered through 0.4 μm syringe filters and loaded onto 10 mL spin columns with 2 mL HisPur cobalt resins (89964, Thermo Scientific). For purification, IMAC columns were first washed with 10 mL protein buffer A and 10 mL of protein buffer A supplemented with 5 mM imidazole. The lysates were then loaded onto IMAC columns and were allowed to bind to the resins for 30 min at 4°C with mixing with a tube rotator. Subsequently, the flowthrough was collected and the bound proteins were eluted in a stepwise manner with 2 × 5 mL of protein buffer A containing increasing concentrations of imidazole (10, 25, 50, 100, 250 and 500 mM). After elution, SDS-PAGE was carried out and relevant fractions containing the protein of interest were pooled and concentrated using Vivaspin 20 filter concentrators (VivaScience) to a volume of <5 mL. The concentrated sample was injected into a pre-flushed sample loop. Size exclusion chromatography was carried out using a Biorad FPLC system with a HiLoad 16/60 Superdex 75 pg column (GE healthcare) pre-equilibrated with protein buffer A. Elution peaks were collected in 1 mL fractions and analysed on SDS-PAGE gels. Relevant peaks were pooled and concentrated using a filter concentrator. The entire purification was performed at 4°C. The final protein concentration was assessed by measuring absorbance at 280 nm on a Nanodrop ND-1000 (Nano-Drop Technologies). The final protein purity was assessed on SDS-PAGE gel (**Fig S2**) and the final protein batches were then aliquoted into smaller fractions, frozen in liquid nitrogen and stored at −80°C.

### SiaC^P^ Preparation by incubating SiaB with SiaC in presence of ATP and size-exclusion chromatography of SiaB/SiaC mixtures

To generate phosphorylated SiaC^P^, SiaB and SiaC proteins were mixed in equal molar ratio and supplemented with 10 mM ATP and 20 mM MgCl_2_. The mixture was incubated at room temperature for 1 h and loaded onto Hiload 16/600 Superdex 200 size exclusion column. SiaC^p^ eluted at approximately 80 mL. Denaturing SDS-PAGE confirmed the size of SiaC^p^, and LC-MS confirmed the phosphorylated state of SiaC^p^. This SiaC^p^ was subsequently used in enzymatic assays using a malachite-green assay. For size-exclusion chromatography SiaB and SiaC mixtures were incubated in the absence and presence of ATP or MgCl_2_ prior to analysis.

### SiaC^P^ production and purification from Pseudomonas aeruginosa cells

For the production of the phosphorylated SiaC^P^ used the Δ*siaA* mutant harbouring the inducible protein production plasmids pJEM[SiaC]. Using glycerol stocks, overnight cultures were inoculated in 20 mL LB medium supplemented with 10 mM phosphate buffer (pH 7.5) and 25 mM glucose and 50 µl/mL kanamycin. The cultures were incubated in 100 mL Erlenmeyer flasks at 30°C with 250 rpm shaking overnight. The following morning, the cultures were used to inoculate fresh 75 mL medium (500 mL Erlenmeyer flask) at an OD_600_ of 0.1. The medium was additionally supplemented with 0.2% rhamnose (w/v) for induction of SiaC production and cultures were incubated at 20°C with 250 rpm shaking. After 16 h of incubation, cells were harvested by centrifugation (9500 × *g*; 5 min; 4°C) and resuspended with 7 mL of ice cold HEPES buffer (20 mM HEPES; pH 7.5) containing 300 mM NaCl, 5 mM imidazole, 5% (v/v) glycerol, a phosphatase inhibitor mix (1× concentrated; Pierce™ Phosphatase Inhibitor Mini Tablets), 0.1 mM Phenylmethylsulfonylfluorid (PMSF); 0.05 mg/mL DNAse, and 0.1 mg mL^-1^ lysozyme. The cell suspension was lysed by sonication at 20% amplitude and a 5 s on/off cycle for 5 min, on ice using a Branson Digital Sonifier 250. The lysate was clarified by centrifugation at 9500 × *g* at 4°C for 10 min. The supernatants were filtered through 0.1 um syringe filters and the SiaC^P^ protein was purified using two GraviTrap™ TALON® columns (GE Healthcare). For purification, TALON® columns were first washed with 4 mL of 20 mM HEPES (pH: 7.5) buffer containing 300 mM NaCl, 5% Glycerol and 500 mM imidazole. Subsequently, 4 mL of dialysis buffer (20 mM HEPES [pH: 7.5], 300 mM NaCl, 5 % glycerol) and 4 mL of equilibration buffer (20 mM HEPES [pH: 7.5], 300 mM NaCl, 5 % glycerol, 5 mM imidazole), 3.5 mL of lysate was loaded onto each of the columns. After the columns were loaded, the bound proteins were washed with 4 mL of HEPES buffer (20 mM HEPES [pH 7.5], 300 mM NaCl, 5 % glycerol) with increasing concentrations of imidazole (10 mM and 25 mM). Finally, the proteins were eluted with 4 mL of HEPES buffer (20 mM HEPES [pH 7.5], 300 mM NaCl, 5 % glycerol) containing 250 mM of imidazole. After elution, the imidazole was diluted from the sample (∼1:10000) and concentrated to ∼ 600 µL using two Vivaspin 6 filter concentrators (VivaScience). The protein concentration of the final sample was assessed by measuring absorbance at 280 nm on a Nanodrop ND-1000 device (Nano-Drop Technologies). The purification process was assessed with SDS-PAGE gel (**Fig S2**) and the final protein batch was aliquoted into smaller fractions, frozen in liquid nitrogen and stored at −80°C.

### Phosphatase assays with purified SiaA-PP2C and fumonisin B_1_

The phosphatase activity of SiaA-PP2C was assessed using para-nitrophenyl phosphate (pNPP) or as the target substrate. Phosphatase activity was determined via the production of a colorimetric para-nitrophenolate product that absorbs strongly at 405 nm. Four times working solutions of the protein (4×: 200 μg/mL; 1×: 50 μg/mL [∼1.49 µM]) were made in water and mixed with equal volume of 4× pNPP phosphatase assay buffer (1×: 20 mM Tris-HCl [pH 7.5], 150 mM NaCl, 20 mM MgCl_2,_ fumonisin B_1_ 0-0.5 mM). Fifty μL of the mixture was then added to each well in 96 well plates, following which pNPP was added to a final concentration of 0-5 mM to make up a final volume of 100 μL per well. Phosphatase activity over time was detected via the measurement of the absorbance at 405 nm using a microtitre plate reader at 37°C (Tecan Infinite M200 Pro). Initial reaction rates were determined using data from 0-10 min (μM/min). Activity of SiaA-PP2C in the presence of fumonisin B1 was calculated as: % Activity = Rate [fumonisin] / Rate [no fumonisin].

### Malachite green phosphate assays and ADP-Glo™ assays

Phosphatase and kinase activities of SiaA and/or SiaB on SiaC and/or SiaC^p^ were determined using the Malachite green phosphate assay kit (Sigma Aldrich) and ADP-Glo™ assay kit (Promega) respectively. 0.5 μM of SiaA and/or SiaB was incubated with 0-5 μM of SiaC or SiaC^p^ in the reaction buffer (20 mM Tris-HCL [pH 7.5], 150 mM NaCl with addition of 0-25 mM ATP and 0-20 mM MnCl_2_ and/or 0-20 mM MgCl_2_) for 1 h at 37°C. Subsequently, the reactions were stopped and the amount of phosphate released in phosphatase reactions or ATP consumed in kinase reactions was determined in accordance to the commercial kit protocols. Briefly, for Malachite green assays, 20 μL of the kit working solution was added to 80 μL of the sample to stop the reaction. The mixture was then incubated at room temperature for 30 min for the generation of a green complex formed between Malachite Green, molybdate and free orthophosphate. Measurements were taken at 620 nm using a microplate reader. Readings were then converted to phosphate measurements using a standard curve. For ADP-Glo™ assays, 20 uL of the ADP-Glo™ reagent was added to 20 uL of the sample to stop the reaction and deplete remaining ATP. If enzyme reactions were carried out in the absence of Mg^2+^, MgCl_2_ was supplemented to the mixture to a final concentration of 20 mM. The mixture was incubated at room temperature for 40 min. A second kinase detection reagent was added to the mixture to convert ADP consumed to ATP and for conversion of the ATP signal to luminescence signals and further incubated for 45 min. Luminescence was measured using a microplate reader and converted to ATP consumed in kinase reaction using a standard curve generated. At least two independent biological replicates with technical duplicates were carried out for each enzymatic reaction.

### Shotgun peptide mass spectrometry of SiaC and evaluation of its phosphorylation state at position threonine 68 as posttranslational modification

Shotgun (fragmentation) peptide mass spectrometry (PMS) was used to evaluate the phosphorylation state of SiaC at threonine residue 68 (T68)[74], for example after reactions with recombinant, purified SiaC as substrate for the phosphatase domain of SiaA (SiaA-PP2C) or kinase SiaB and ATP. For such assays, 20 µg of purified SiaC^P^ or SiaC was incubated in the absence or presence of 20 µg of purified SiaA or SiaB for 2 h at 30°C in suitable assay solution (see above). Twenty µL were used for the separation of the proteins within the sample using a 15% SDS-PAGE gel [104]. The gel was stained with Coomassie Blue [105] and the SiaC protein band (17 kDa) was excised from the SDS-PAGE gel, subjected to tryptic digest and the peptides were analyzed by PMS at the Proteomics Centre of University of Konstanz, as described previously with minor modifications [106–108]. Here, each sample was analyzed twice on a Orbitrap Fusion with EASY-nLC 1200 (Thermo Fisher Scientific) and Tandem mass spectra were searched against an appropriate protein database (retrieved from IMG) using Mascot (Matrix Science) and Proteome Discoverer V1.3 (Thermo Fisher Scientific) with “Trypsin” enzyme cleavage, static cysteine alkylation by chloroacetamide, and variable methionine oxidation, serine and threonine phosphorylation.

For analysis, the relative abundance of precursor ions was determined for the peptide harboring the T68 in a non-phosphorylated state (parental peptide [LLYLN**T**SSIK]) and for the peptide with corresponding mass-shifts indicative of its phosphorylation (monoisotopic mass-change, +79.96633 Da; average mass-change +79.9799 Da).

### Crystallization, data collection and structure solution

All crystals were cryo-protected with 20% glycerol and frozen in liquid nitrogen and X-ray diffraction data were collected at beamline PROXIMA 2A of SOLEIL, France. Purified SiaA-PP2C (386-663) and SeMet SiaA-PP2C (386-663) proteins were concentrated to 25 mg/mL and 24 mg/mL respectively in 20 mM Tris-HCl PH 8, 500 mM NaCl and 1 mM DTT for screening of crystallization conditions. The best SiaA-PP2C native and SeMet derivative crystals were obtained from Morpheus A11 and A6 of Molecular Dimension, respectively. SAD phasing and initial model building was processed with AutoSol from the Phenix package [109] followed by structure refinement with buster [110] and PDB deposition (6K4E). The SeMet derivative crystal of SiaC was first obtained in condition Morpheus A9 of Molecular Dimensions at resolution 2.9 Å. To improve resolution, the His-tag of SiaA protein was removed by TEV cleavage, prior to the native crystal screen. A native SiaC crystal was obtained in 0.1 M Phosphate/citrate, pH 4.2, 40% v/v PEG 300 at 1.7 Å. SAD phasing and initial model building was processed with AutoSol from the Phenix package, followed by structure refinement with Phenix Refine and PDB deposition (6K4F).

### Homology modelling, molecular docking and molecular dynamics (MD) simulations

The homology model of SiaB was generated with the I-TASSER online Server by specifying the amino acid sequence of SiaB. The homology model of SiaA-PP2C* was generated with SWISS-MODEL by providing the amino acid sequence of SiaA-PP2C* and specifying the SiaA-PP2C crystal structure as template.

SiaC^P^ model was generated by substituting T68 with pT68 using psfgen in VMD [111]. The model was then subjected to conjugate gradient energy minimization for 10000 steps using NAMD 2.11 [112]. SiaC^P^ was docked to the SiaA dimer by ZDOCK 3.0.2 [113] by specifying pT68 as interacting residues. Among the top 10 complexes, complex 2 was chosen as the best model because the pT68 was closest to the bridging water.

SiaC^P^-SiaA and SiaC-SiaA dimers were subjected to all-atom, explicit solvent MD simulation using NAMD 2.11. Each dimer model was simulated in a water box, where the minimal distance between the solute and the box boundary was 15 Å along all three axes. The charges of the solvated system were neutralized with counter-ions, and the ionic strength of the solvent was set to 150 mM NaCl using VMD. The fully-solvated system was subjected to conjugate-gradient minimization for 10,000 steps, subsequently heated to 310 K, and a 10 ns equilibration with protein backbone atoms constrained using a harmonic potential of the form U(x) = k (x-x_ref_)^2^, where k is 1 kcal/mol Å^-2^ and x_ref_ is the initial atom coordinates. Finally, 60 ns production simulations were performed without constraints. All simulations were performed under the NPT ensemble assuming the CHARMM36 force field for the protein and the TIP3P model for water molecules. PDB files of the homology models and docking results used for the study are provided as zipped Supplementary files.

### Statistical analysis

When needed, experiments with n ≥ 3 were analysed using a two-sided students *t*-tests (α = 0.05) with *p*-values < 0.05 interpreted as being significantly different.

## Acknowledgements

The authors further declare that the research was conducted in the absence of any commercial or financial relationships that could be construed as a potential conflict of interest. Janosch Klebensberger would like to thank Prof. Bernhard Hauer for his continuous support. We also acknowledge beam time allocation at the SOLEIL synchrotron. The MD simulations for this article were performed on ASPIRE 1 of the National Supercomputing Centre, Singapore (https://www.nscc.sg).

## Supporting Information

**Fig S1:**
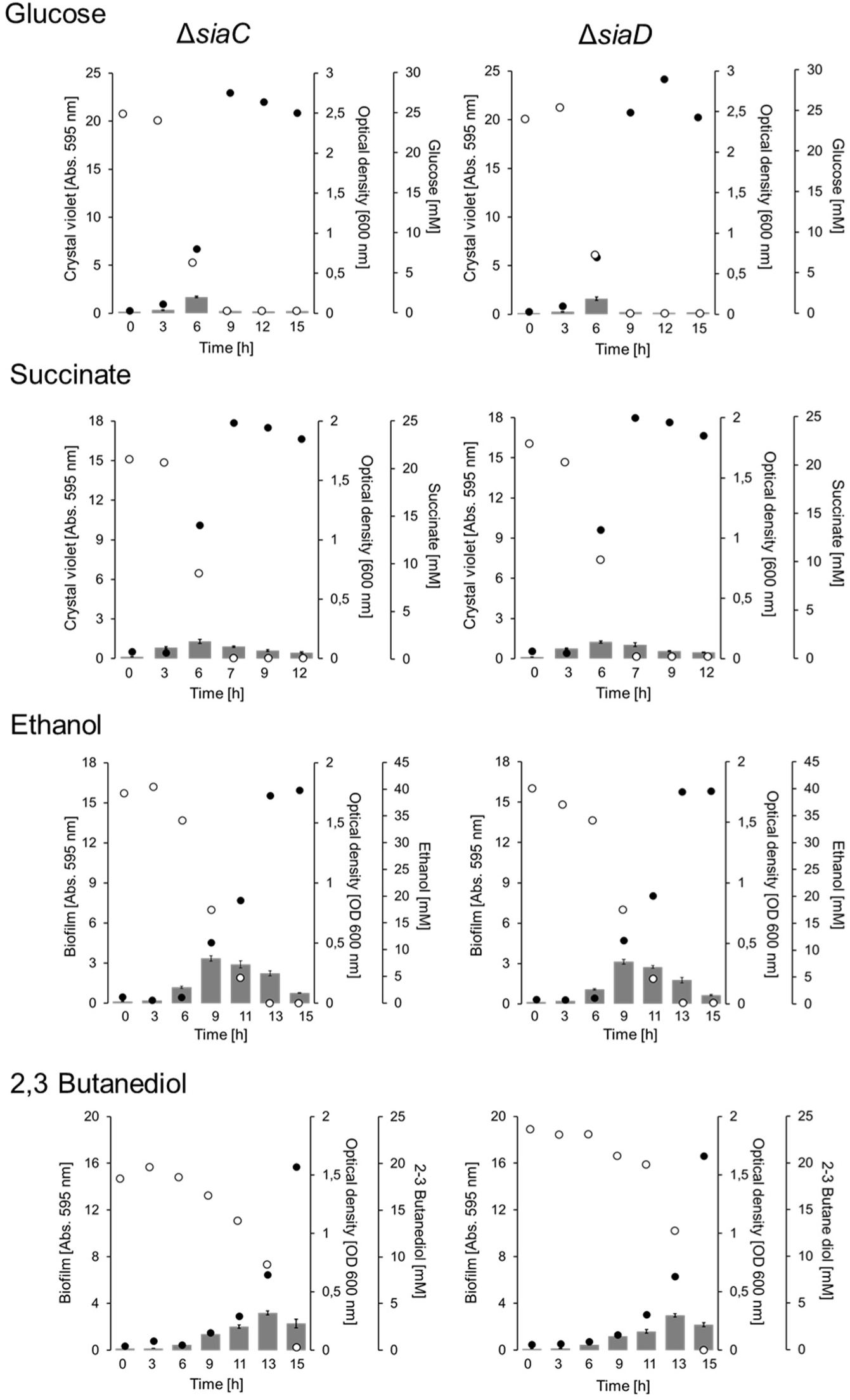
Growth of wild type and the Δ*siaC* and Δ*sia*D strains in M9 medium with 22 mM glucose, 20 mM succinate, 40 mM ethanol or 20 mM 2,3 butanediol at 30°C and 200 rpm shaking. Individual cell-culture treated, 24 well microtiter plates were used to quantify attached biofilms (*grey bars*), OD_600_ of supernatants (*black dots*), and substrate concentrations (*open circles*) at various time points. Attached biofilms were quantified by a modified crystal violet (CV) staining and error bars represent the standard deviation of four biological replicates (n = 4). OD_600_ of supernatants and substrate concentrations represent the mean value from the same quadruplicates but quantified from pooled (1:1:1:1 [v/v/v/v]) samples using a photometer, the GO assay kit, and specific HPLC methods (see Material and Method for details).

**Fig S2:**
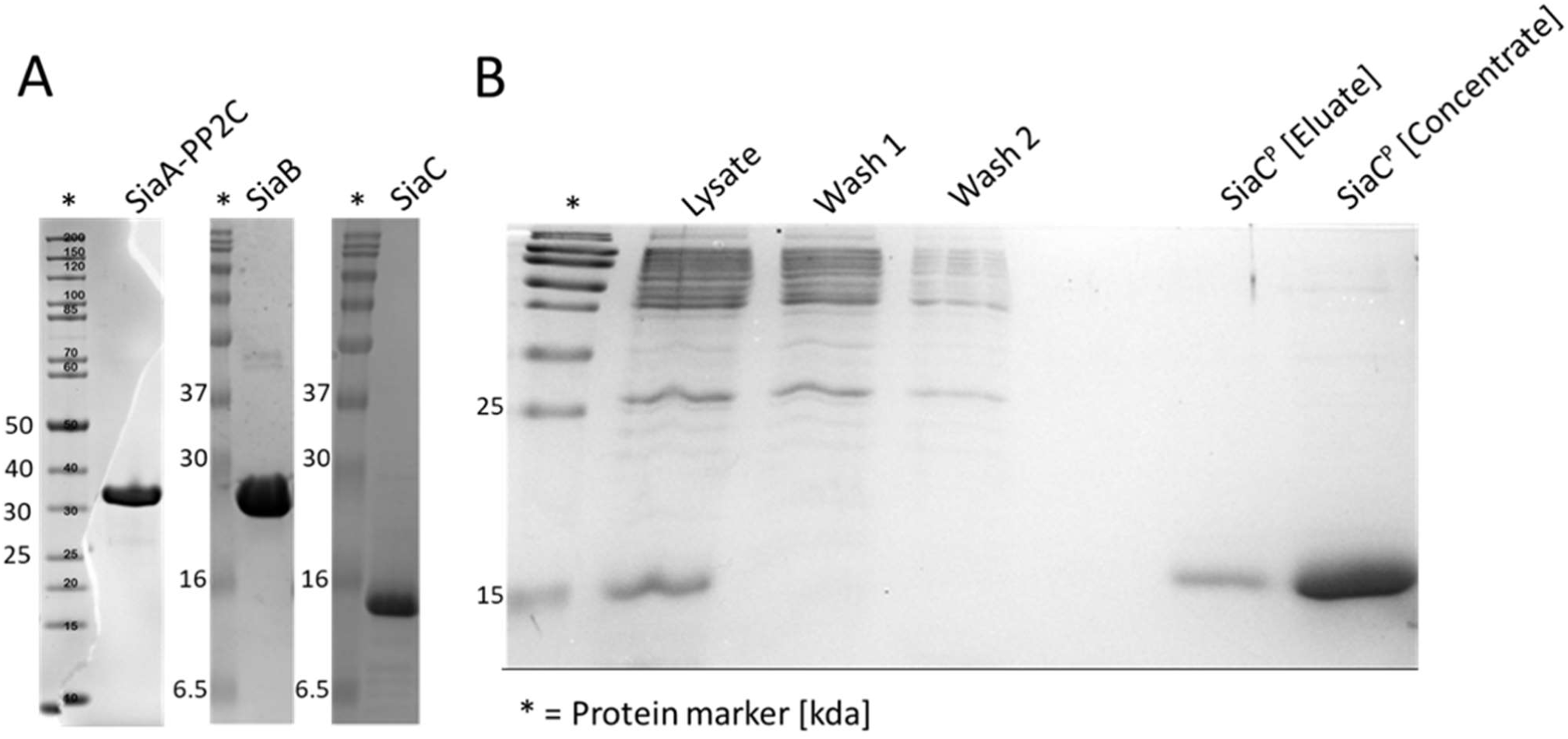
SDS-PAGE analysis of the purified SiaA-PP2C, SiaB, SiaC and SiaAPP2C* protein samples after purification. A) The purified SiaA phosphatase domain (amino acids 386-663 of the SiaA protein sequence; Genbank ID: NP_248862), the SiaB (Genbank-ID: 248862) and SiaC (Genbank-ID: NP_248860) proteins produced in *E. coli* after affinity chromatography and subsequent gel filtration. B) The purified SiaC protein (Genbank-ID: NP_248860) produced in the Δ*siaA* mutant (SiaC^P^) after affinity chromatography (eluate) and subsequent protein concentration.

**Fig S3:**
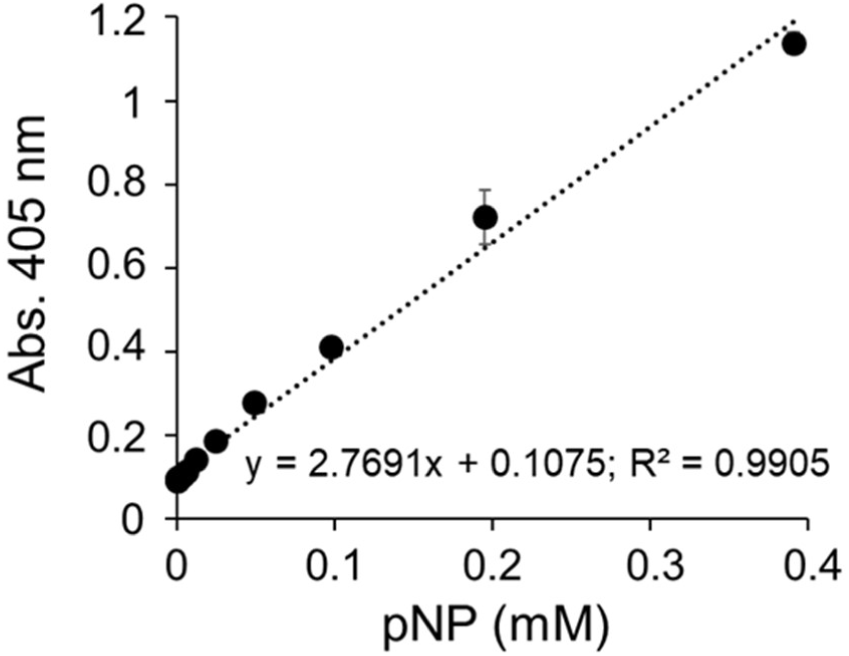
Standard calibration curve of p-nitrophenolate (pNP) concentrations against absorbance (Abs. 405 nm) in phosphatase assay buffer (150 mM NaCl, 100 mM Tris-HCl pH 7.5).

**Fig S4:**
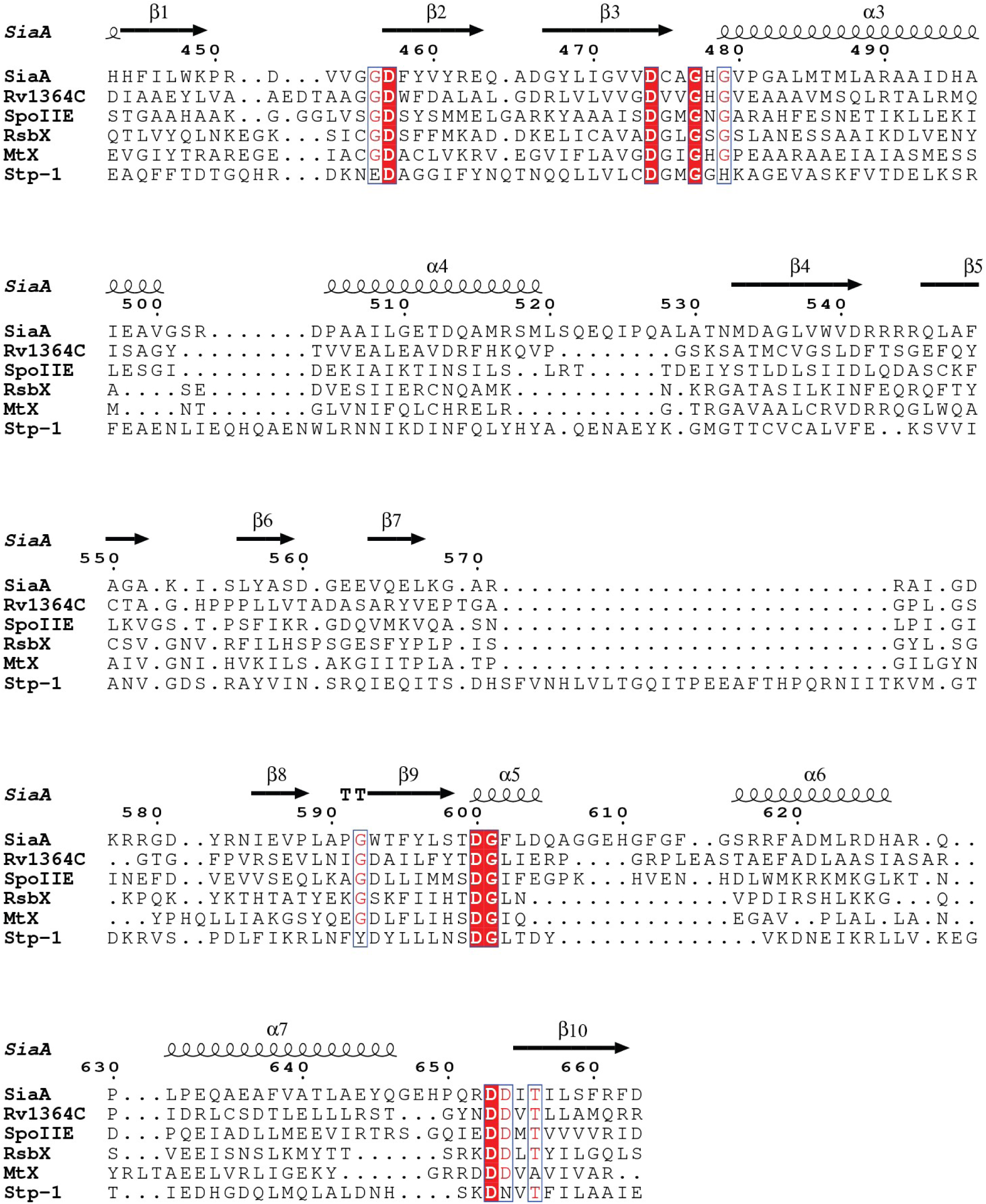
Structure-based sequence alignment of SiaA-PP2C against other PP2C-type phosphatases, identified by an automated search using the Dali server (ekhidna2.biocenter.helsinki.fi/dali) of 3D structures. Conserved residues are highlighted in red. Secondary structure features and residue number of SiaA-PP2C are labelled on top of the alignment.

**Table S1:**
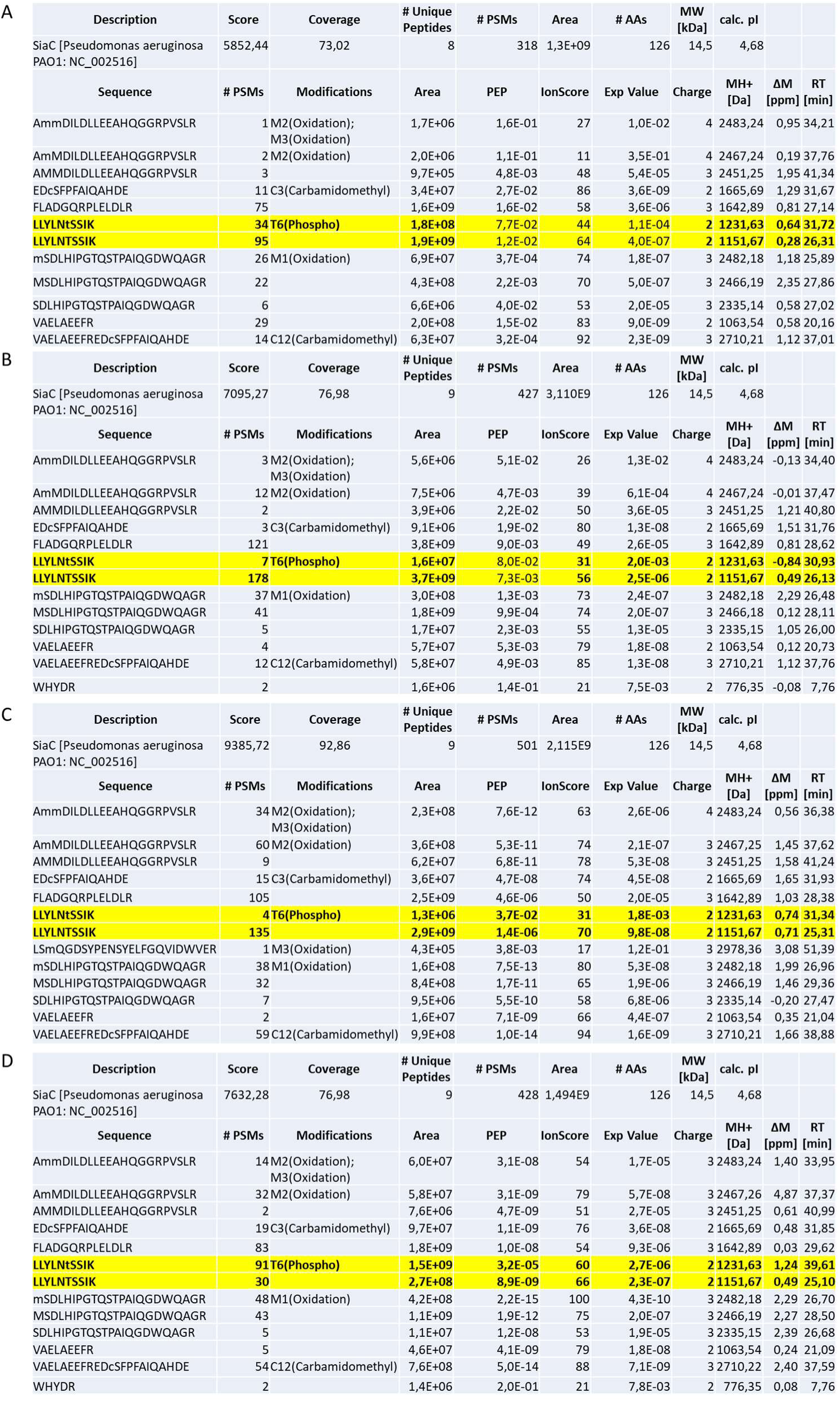
Summary of peptide fingerprinting-mass analysis of the SiaC protein after various incubations. Analysis of SiaC^P^ (purified from *P. aeruginosa*) in the absence **(A)** and presence of the phosphatase domain of SiaA (SiaA-PP2C) **(B)** after incubation for 2 h at 30°C in phosphatase buffer. SiaC (purified from *E. coli*) and SiaB incubated in the absence **(C)** and presence **(D)** of ATP after incubation for 2 h at 30°C in kinase buffer. Subsequently, the SiaC protein was separated by SDS-PAGE; the SiaC-protein band was cut out of the gel and analysed by PF-MS.

**Table S2:** X-ray diffraction data collection, solution and statistics of the native and SeMet crystals of SiaA-PP2C and SiaC.

|  | SiaA Native | SiaA SeMet | SiaC Native | SiaC SeMet |
| --- | --- | --- | --- | --- |
| **Wavelength (Å)** | 0.9764 | 0.979 | 0.9801 | 0.9766 |
| **Resolution range (Å)** | 49.81 - 2.094<br>(2.169 - 2.094) | 27.35 - 2.49<br>(2.579 - 2.49) | 57.96 - 1.735<br>(1.797 - 1.735) | 38.74 - 2.94<br>(3.045 - 2.94) |
| **Space group** | P 21 21 21 | P 21 21 21 | I 2 2 2 | P 31 2 1 |
| **Unit cell (Å)** | 51.17 110.48<br>111.6 90 90 90 | 51.354 109.416<br>111.46 90 90 90 | 35.4 72.8 95.8<br>90 90 90 | 77.482 77.482<br>85.02 90 90 120 |
| **Total reflections** | 205960 (19009) | 606993 (59041) | 116718 (7271) | 130598 (12091) |
| **Unique reflections** | 37739 (3471) | 22684 (2229) | 13113 (1176) | 6550 (603) |
| **Multiplicity** | 5.5 (5.5) | 26.8 (26.5) | 8.9 (6.2) | 19.9 (19.9) |
| **Completeness (%)** | 99.18 (93.11) | 99.85 (99.82) | 98.94 (90.81) | 99.53 (96.20) |
| **Mean I/sigma (I)** | 7.20 (0.99) | 20.17 (1.65) | 12.05 (0.90) | 21.22 (1.56) |
| **Wilson B-factor (Å<sup>2</sup>)** | 41.65 | 66.72 | 34.21 | 97.73 |
| **R-merge** | 0.1603 (1.432) | 0.1411 (1.813) | 0.09846<br>(1.129) | 0.1306 (1.444) |
| **CC1/2** | 0.993 (0.461) | 0.998 (0.752) | 0.998 (0.62) | 0.999 (0.784) |
| **R-work** | 0.2177 (0.3346) | 0.2055 (0.2768) | 0.1972<br>(0.3956) | 0.2122 (0.3450) |
| **R-free** | 0.2522 (0.3782) | 0.2542 (0.3665) | 0.2348<br>(0.4305) | 0.2812 (0.4126) |
| **RMS (bonds) (Å)** | 0.014 | 0.014 | 0.008 | 0.015 |
| **RMS (angles) (°)** | 1.71 | 1.77 | 1.19 | 1.82 |
| **Ramachandran outliers (%)** | 0.00 | 0.00 | 0.00 | 0.41 |
| **Rotamer outliers (%)** | 3.73 | 4.07 | 0.00 | 12.50 |
| **Clashscore** | 2.80 | 3.23 | 2.65 | 8.62 |
| **Average B-factor (Å<sup>2</sup>)** | 53.79 | 71.23 | 43.58 | 106.10 |

**Table S3.** Strains and plasmids used in the study.

| Strains | Relevant features | Reference |
| --- | --- | --- |
| ***P. aeruginosa*** |  |  |
| PAO1 | Wild-type *Pseudomonas aeruginosa* | Holloway Collection |
| $\Delta siaA$ | PAO1 with a 6 bp deletion in *siaA* | [1] |
| $\Delta siaB$ | PAO1 with a markerless *loxP*-site insertion in *siaB* (PA0171) | This study |
| $\Delta siaC$ | PAO1 with a markerless *loxP*-site insertion in *siaC* (PA0170) | This study |
| $\Delta siaD$ | PAO1 with a markerless *res*-site insertion in *siaD* (KO0169) | [1] |
| PW1292 | MPAO1 with transposon phoAwp091G03 inserted at bp 311 of the coding region of the *siaB* gene[ | [2] |
| PW1290 | MPAO1 with transposon phoAwp09q2A10 inserted at bp 181 of the coding region of the *siaC* gene | [2] |
| ***E. coli*** |  |  |
| DH5α | *fhuA2 lac(del)U169 phoA glnV44 Φ80' lacZ(del)M15 gyrA96 recA1 relA1 endA1 thi-1 hsdR17* | [3] |
| S17-λpir | Tp<sup>r</sup> Sm<sup>r</sup> *recA*, *thi*, *pro*, *hsdR*-M<sup>+</sup>RP4: 2-Tc:Mu: Km<sup>r</sup> Tn7 λpir | [4] |
| BL21(DE3) Rosetta T1R | *E. coli* str. B F<sup>-</sup> *ompT gal dcm lon hsdS<sub>B</sub>(r<sub>B</sub><sup>-</sup>m<sub>B</sub><sup>-</sup>) $\lambda$(DE3 [*lacI lacUV5-T7p07 ind1 sam7 nin5*]) [*malB*<sup>+</sup>]<sub>K-12</sub>($\lambda^S$) *tonA** | NTU Protein Production Plattform |
| **Plasmids** |  |  |
| pCR2.1 | TOPO TA cloning plasmid, Amp<sup>r</sup>, Kan<sup>r</sup> | Invitrogen |
| pBBR | Broad host range expression plasmid pBBR1MCS-5, Gm<sup>r</sup> | [5] |
| pJEM1 | Rhamnose inducible broad host range expression plasmid, Kan<sup>r</sup> | [6] |
| pNIC28-BSA4 | Protein production plasmid, N-terminal His6, TEV-cleavable MHHHHHHSSGVDLG TENLYFQ*SM, Kan<sup>r</sup> | [7] |
| pNIC-CTHF | Protein production plasmid, Cleavable C-terminal His6-FLAG tag, Kan<sup>r</sup> | [7] |
| pNIC[SiaA] | pNIC28-BSA4 harbouring the partial *siaA* gene (C-terminal phosphatase domain from amino acid 386-663) for the production of a N-terminal His6-TEV-SiaA allele (SiaA-PP2C). | This study |
| pNIC[SiaB] | pNIC28-BSA4 for the production of a N-terminal His6-TEC SiaB allele | This study |
| pNIC[SiaC] | pNIC28-BSA4 for the production of a N-terminal His6-TEV-SiaC allele | This study |
| pJEM[SiaC] | pJEM1 harbouring the *siaC* gene for the production of a N-terminal His6-TEV-SiaC allele in *Pseudomonas aeruginosa* strains | This study |
| pCre1 | Plasmid for the expression of Cre recombinase | [8] |
| **Phage** |  |  |
| E79tv2 | Generalised transducing phage | [9] |

---

**Table S4:**
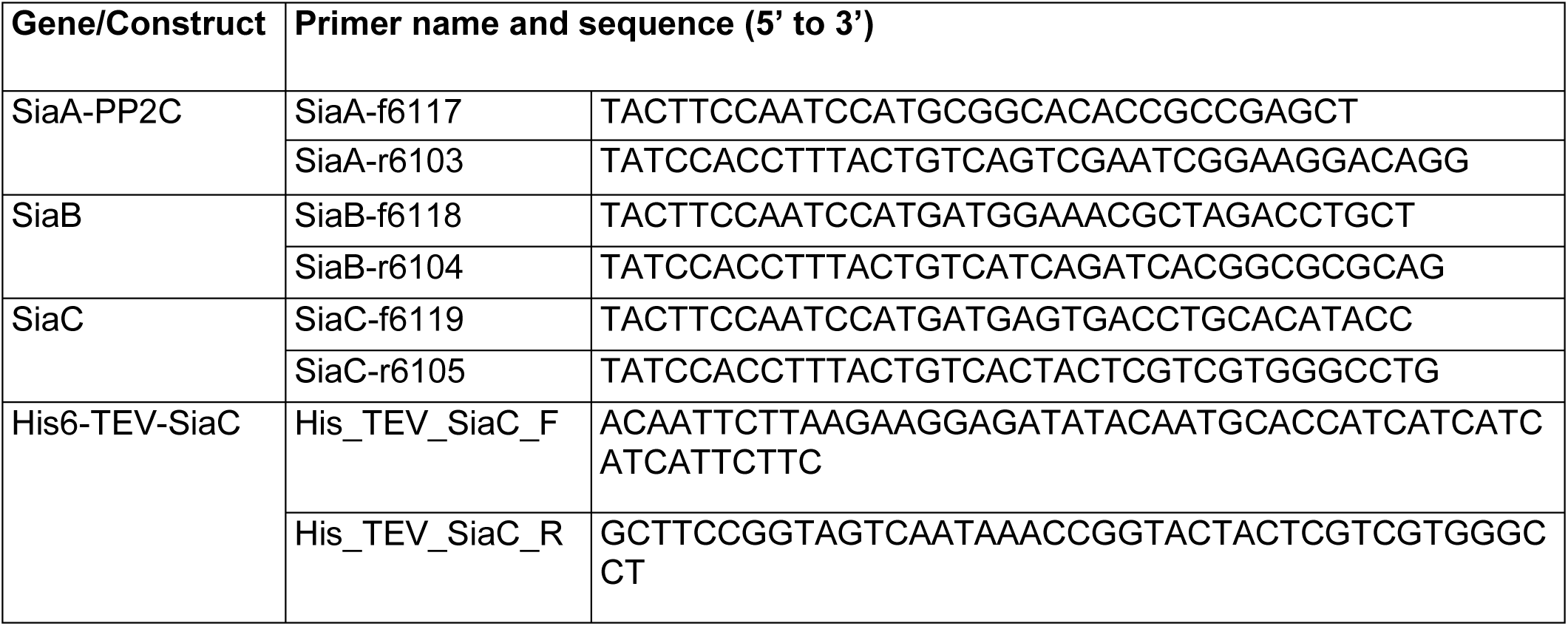
Primers used in the study.

## Supplementary zip-file

1. SiaB model_I-Tasser
2. SiaC-PhosT68 model_Energy minimised
3. SiaAC complex_MD simulation_60ns2
4. SiaAC-T68P complex_MD simulation_60ns2
5. SiaA-PP2C_Mutant allele_homology model

